# Spatial population dynamics of bacterial colonies with social antibiotic resistance

**DOI:** 10.1101/2024.08.21.608973

**Authors:** Marlis K. Denk-Lobnig, Kevin B. Wood

## Abstract

Bacteria frequently inhabit surface-attached communities where rich “social” interactions can significantly alter their population-level behavior, including their response to antibiotics. Understanding these collective effects in spatially heterogeneous communities is an ongoing challenge. Here, we investigated the spatial organization that emerges from antibiotic exposure in initially randomly distributed communities containing antibiotic-resistant and -sensitive strains of *E. faecalis*, an opportunistic pathogen. We identified that a range of complex spatial structures emerged in the population homeland—the inoculated region that microbes inhabit prior to range expansion—, which depended on initial colony composition and antibiotic concentration. We found that these arrangements were explained by cooperative interactions between resistant and sensitive subpopulations with a variable spatial scale, the result of dynamic zones of protection afforded to sensitive cells by growing populations of enzyme-producing resistant neighbors. Using a combination of experiments and mathematical models, we explored the complex spatiotemporal interaction dynamics that create these patterns, and predicted spatial arrangements of sensitive and resistant subpopulations under new conditions. We illustrated how spatial population dynamics in the homeland affect subsequent range expansion, both because they modulate the composition of the initial expanding front, and through long-range cooperation between the homeland and the expanding region. Finally, we showed that these spatial constraints resulted in populations whose size and composition differed markedly from matched populations in well-stirred (planktonic) cultures. These findings underscore the importance of spatial structure and cooperation, long-studied features in theoretical ecology, for determining the fate of bacterial communities under antibiotic exposure.

**Significance:** Interactions between bacteria are common, particularly in the crowded surface-associated communities that occur anywhere from natural ecosystems to the human body to medical devices. Antibiotic resistance can be influenced by these “social” interactions, making it difficult to predict how spatial communities respond to antibiotic. Here, we show that complex spatial arrangements emerge when initially randomly distributed populations of antibiotic-resistant and -sensitive *E. faecalis*, a microbial pathogen, are exposed to antibiotic. Using mathematical models and experiments, we show how local competition and dynamic-range cross-protection drive pattern formation. As a result, these spatially structured populations respond differently to antibiotics than well-mixed communities. Our findings elucidate how “social” antibiotic resistance affects spatially structured bacterial communities, a step towards predicting and controlling resistance.

## Introduction

### Spatial structure as an integral feature of life

Spatial organization is a fundamental property of life across scales; how molecules and cells organize into complex, coordinated organisms and ecosystems is an astounding and counter- intuitive process. Whereas bacteria are often thought of as individual cells swimming in a well-mixed liquid environment, the majority of bacterial life exists as surface-attached, crowded and spatially structured communities, such as biofilms ^1–3^. In these densely packed environments, interactions between organisms abound and collective behaviors emerge at population and community scales ^4–6^. Recent work on collective behaviors of bacteria has made it increasingly clear that appreciating the “social lives” of bacteria in these communities is essential to better predict and potentially control their behaviors, whether in natural ecosystems, healthy microbiomes or infections ^7–10^.

### Bacterial resistance to antibiotics is a “social” phenomenon

Resistance against antibiotic compounds is increasingly understood as a collective, eco- evolutionary process ^11–14^. Antibiotic resistance is a rising threat to global public health, rapidly reducing treatment options for bacterial infections ^15,16^. Developing new antibiotics is an arduous process ^17^, which makes it critical to better understand how bacterial populations interact with existing antibiotics, in hopes of prolonging their use and reducing the spread of resistance. One promising approach calls for shifting focus from molecular-level mechanisms to (microbial) population-level ecological and evolutionary drivers of antibiotic resistance ^18–21^.

Antibiotic resistance frequently involves interactions between bacteria, and those interactions affect the establishment and spread of resistance in a bacterial community ^22^. Interactions are often mediated by direct modulation of antibiotic concentrations—for example, cells with increased efflux activity increase local extracellular antibiotic concentrations ^23^, while cell producing antibiotic-degrading enzymes reduce it ^24–26^. β-lactamases, enzymes that degrade β- lactam antibiotics such as penicillin or ampicillin and have been found in a wide array of bacterial species, represent a classic mechanism of collective resistance ^25–28^. In homogeneous β-lactamase expressing populations, antibiotic degradation leads to an inoculum effect, where denser initial cultures rapidly detoxify their environment and thus display improved survival compared to lower-density cultures ^29^. In communities that include subpopulations with and without β-lactamase, non-producing bacteria can act as cheaters, reaping the benefit of the detoxification without contributing to it ^30–32^. Interactions between resistant and non-resistant subpopulations that aren’t directly related to the antibiotic, such as competition for nutrients and space, active war-fare and mutualistic exchange of nutrients, can also shift the balance between subpopulations and thus affect community resistance ^33–35^. For example, antibiotic-sensitive cells can compete for nutrients with antibiotic-resistant bacteria, reducing the overall resistance levels of the community or even contributing to community collapse ^19,36–38^. The interplay of such ecological interactions can lead to counterintuitive emergent behaviors and resistance levels at the population and community scales that are not obvious from molecular or single-cell resistance mechanisms. Predicting these collective behaviors is an ongoing challenge to understanding and strategically combatting antibiotic resistance at the population level.

### Social interactions are influenced by spatial organization

Spatial structure is known to play important roles in eco-evolutionary processes generally, both in microbial systems and in macroscopic scale ecosystems of plants and animals, because it shapes when and where different ecosystem components interact ^6,39–41^. In an agar colony or biofilm, bacteria at different spatial positions may experience different environmental conditions, from nutrient and oxygen concentrations to physical pressures ^42,43^. This heterogeneity can be important for ecological interactions, and those interactions can inversely drive emergent spatial structure. Bacterial communities across various systems have been known to form collective spatial patterns through local or spatially heterogeneous interactions ^44–49^. Spatial heterogeneity can promote cooperative secretion of public goods if cooperators can physically segregate from cheaters to retain benefits of production within their group ^50–54^. It can also lead to stochastic outcomes of population growth, as the initial positioning or slight differences in growth timing can decide whether a subpopulation or lineage is able to expand ^55,56^. These social interactions occur at a large range of distinct length scales, from “short-range” metabolic interactions on the micron scale to long-range protective effects in some antibiotic-degrading populations ^35,57–59^. The biophysical determinants of these length scales are not fully understood, as it is difficult to disentangle the complex interplay between local population growth and the spatiotemporal gradients of antibiotic concentration that modulate, and are modulated by, that growth. Thus, the relationship between spatial structure and the dynamics of ecologically interacting subpopulations, especially in the context of antibiotic resistance, is not fully understood.

### How do social populations of bacteria respond to antibiotic in a spatially organized context?

In this work, we investigated the relationship between spatial structure and “social” antibiotic (β-lactam) resistance in populations of *Enterococcus (E.) faecalis* growing on 2D agar surfaces.

*E. faecalis* is a Gram-positive opportunistic pathogen that frequently exhibits and easily acquires antibiotic resistance ^60^. *E. faecalis* is a useful model organism for this question because 1) it is immotile, 2) it has a round shape that is not particularly sensitive to β-lactam concentration ^61^, and 3) β-lactamase produced by resistant cells is not excreted into the extracellular space but remains cell-associated ^62,63^. This strips away potential complexities associated with cell motility, cell shape or filamentation, and diffusive dynamics of the enzyme itself ^37,64,65^.

While a significant body of research has focused on range expansions, where ecological dynamics shape the leading fronts at the edge of expanding microbial colonies ^35,52,66–68^, here we instead focused on colony growth in the so-called “homeland”, where the dilute inoculum gives rise to multiple clonal founder populations that potentially compete, and cooperate, to fill the colony before range expansion even begins. More specifically, we examined the spatial organization of the homeland in mixed colonies comprised of β-lactamase producing resistant cells and antibiotic-sensitive “cheaters”, using a combination of confocal microscopy and simple biophysics models. By tuning antibiotic concentration and the composition and density of the seeding population, we identified a wide range of spatial arrangements of antibiotic-resistant and sensitive bacteria that can be explained by coupling cell proliferation and death to a simple combination of ecological interactions between the two subpopulations: local spatial competition and variable-range cross-protection that is driven by reaction-diffusion dynamics of antibiotic degradation. We then showed that these spatially structured population dynamics in the homeland have important consequences for community composition and subsequent range expansion.

## Results

### The homeland of *E. faecalis* cooperative resistant and sensitive colonies forms a wide range of spatial arrangements under ampicillin exposure

To investigate the spatial organization of antibiotic-resistant *E. faecalis* populations, we created mixed communities comprised of an ampicillin-resistant *E. faecalis* og1rf strain, which constitutively expresses a plasmid-encoded β-lactamase as well as a green fluorescence marker (“resistant” strain, green), and an ampicillin-sensitive strain harboring an enzyme-free variant of the plasmid expressing a magenta marker (“sensitive” strain, magenta) ^57,69^. For each experiment, we placed a well-mixed 2 µl droplet with specified cell density and composition (ratio of resistant and sensitive bacteria) onto an agar plate containing a defined ampicillin concentration, and allowed the cells to grow overnight at 37°C (Figure 1). To image the colonies, we designed a simple 3D-printed adaptor that allowed us to directly image the unperturbed homeland using confocal microscopy. Using this setup, we discovered that the resistant and sensitive populations grow into complex and varied spatial arrangements (Figure 1B, Figures S1-S3). The composition and spatial organization were largely constant along the entire depth of the colony (Figure S4), suggesting that the dynamics take place in an effectively two-dimensional environment. We found that sensitive cells were able to survive in mixed communities at otherwise lethal antibiotic concentrations (Figure S5), consistent with cooperative resistance ^70^. The spatial structures emerging from this process strongly depended on the ampicillin concentration as well as the initial density and resistant fraction of the colony. The scale of protection at times appeared long-range, with sensitive cells surviving even when separated from resistant cells by tens or even hundreds of cell lengths (∼1 µm).

**Figure 1:**
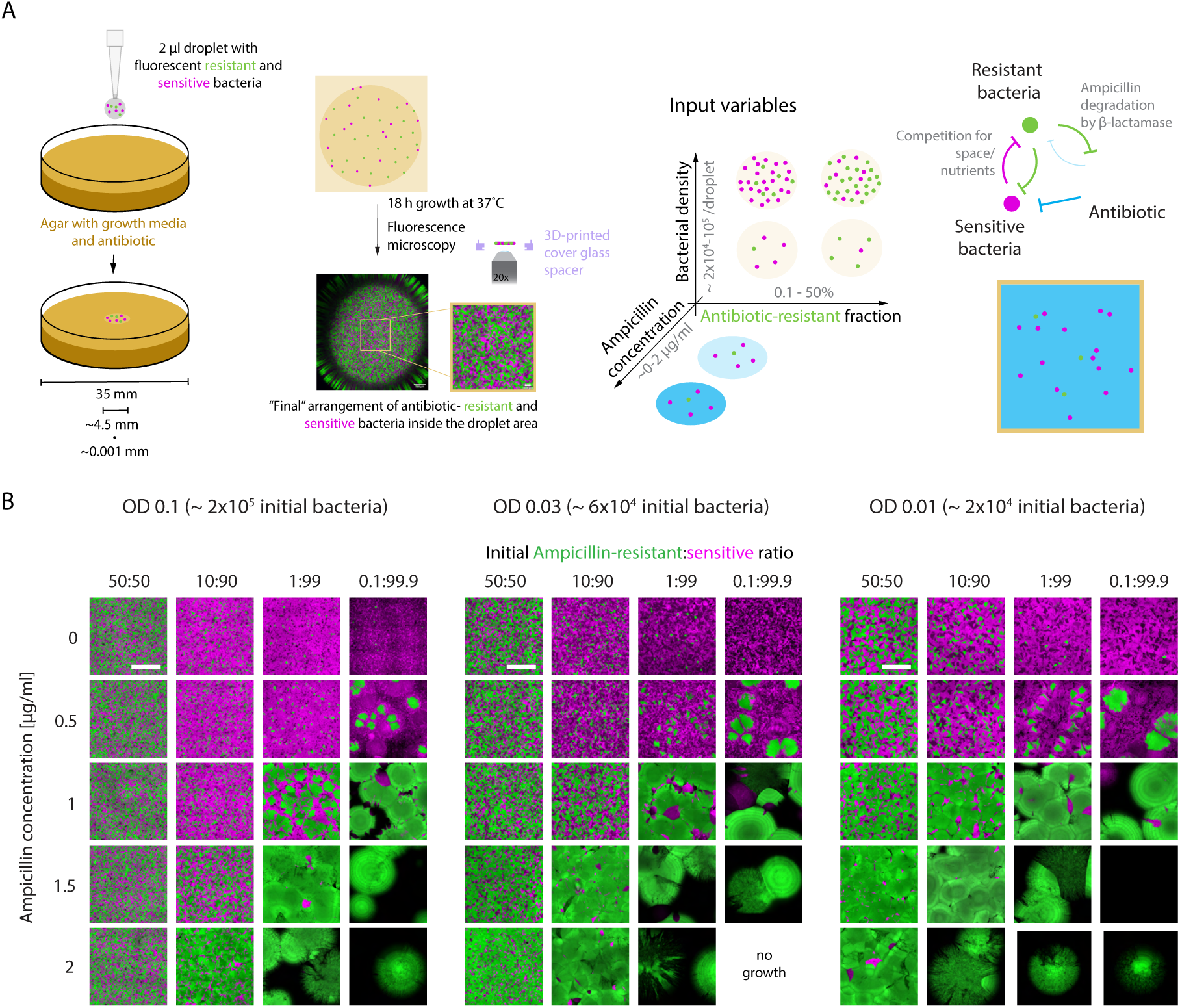
Growth of β-lactamase positive and negative *E. faecalis* subpopulations forms a variety of spatial arrangements in the homeland of agar colonies exposed to the antibiotic ampicillin. (A) Experimental setup to investigate homeland spatial organization. Left: A mixture of defined density and ratio of fluorescently labeled ampicillin-resistant *E. faecalis* bacteria expressing β-lactamase and ampicillin sensitive *E. faecalis* bacteria is added in a 2µl droplet to an agar plate containing growth media (Brain Heart Infusion), ampicillin, and spectinomycin to select for the fluorescent plasmids. Approximate scales of the plate diameter, droplet diameter and cell diameter are shown. Center left: The seeded colony grows overnight at optimal growth temperature, and is then imaged on a confocal fluorescence microscope, using a cover glass spacer to leave the colony at the agar-air interface unperturbed. The spatial arrangement inside the seeded colony area (the homeland) is analyzed. Center right: Overall density and ampicillin-resistant fraction of the colony, as well as ampicillin concentration were the input variables that were varied across different initial conditions to test their impact on spatial pattern formation. Right: Ampicillin-resistant and sensitive subpopulations are expected to have the following ecological interactions: Competition for space or nutrients within the colony area, and ampicillin-induced death that is alleviated by degradation of antibiotic by the β-lactamase expressed by resistant bacteria. As these immotile bacteria maintain relatively fixed positions in the spatially structured colony apart from cell-division and death processes, these ecological interactions are expected to be heterogeneous and neighborhood-dependent. Resistant cells are shown in green, sensitive cells in magenta and antibiotic concentration is encoded in blue. (B) Resistant and sensitive *E. faecalis* subpopulations form a diverse array of spatial patterns on agar. Representative images (maximum intensity z-projections) of spatial homeland organization of ampicillin-resistant (green, fluorophore mDasher) and ampicillin-sensitive (magenta, fluorophore mRudolph) bacteria after overnight growth. Scale bars = 500 µm; unless otherwise noted, all images in the same panel are at the same scale. “Rings” of varied intensity of the green signal at high ampicillin concentrations and low initial resistant fractions are imaging artifacts arising from the maximum intensity projection in combination with non-uniform illumination across angled surfaces. No growth occurred in some cases with low resistant density and high ampicillin concentration because the β-lactamase resistance mechanism is cooperative and thus at low densities, individual resistant cells were not always sufficiently resistant to survive at 1.5 or 2 µg/ml ampicillin.

### Final population composition depends on initial properties of the colony and its environment

To dissect the pattern formation process between sensitive and cooperative resistant subpopulations, we first described how quantitative features of these spatial patterns are influenced by the initial conditions in each colony. To start, we quantified final population composition by using a machine-learning pixel-classification method ^71^ to estimate the global resistant fraction in the homeland. Under ampicillin-free conditions, the final resistant fraction matched or slightly exceeded the initial resistant fraction, suggesting that the resistant strain did not experience significant fitness costs in the absence of antibiotic (Figure 2A).

**Figure 2:**
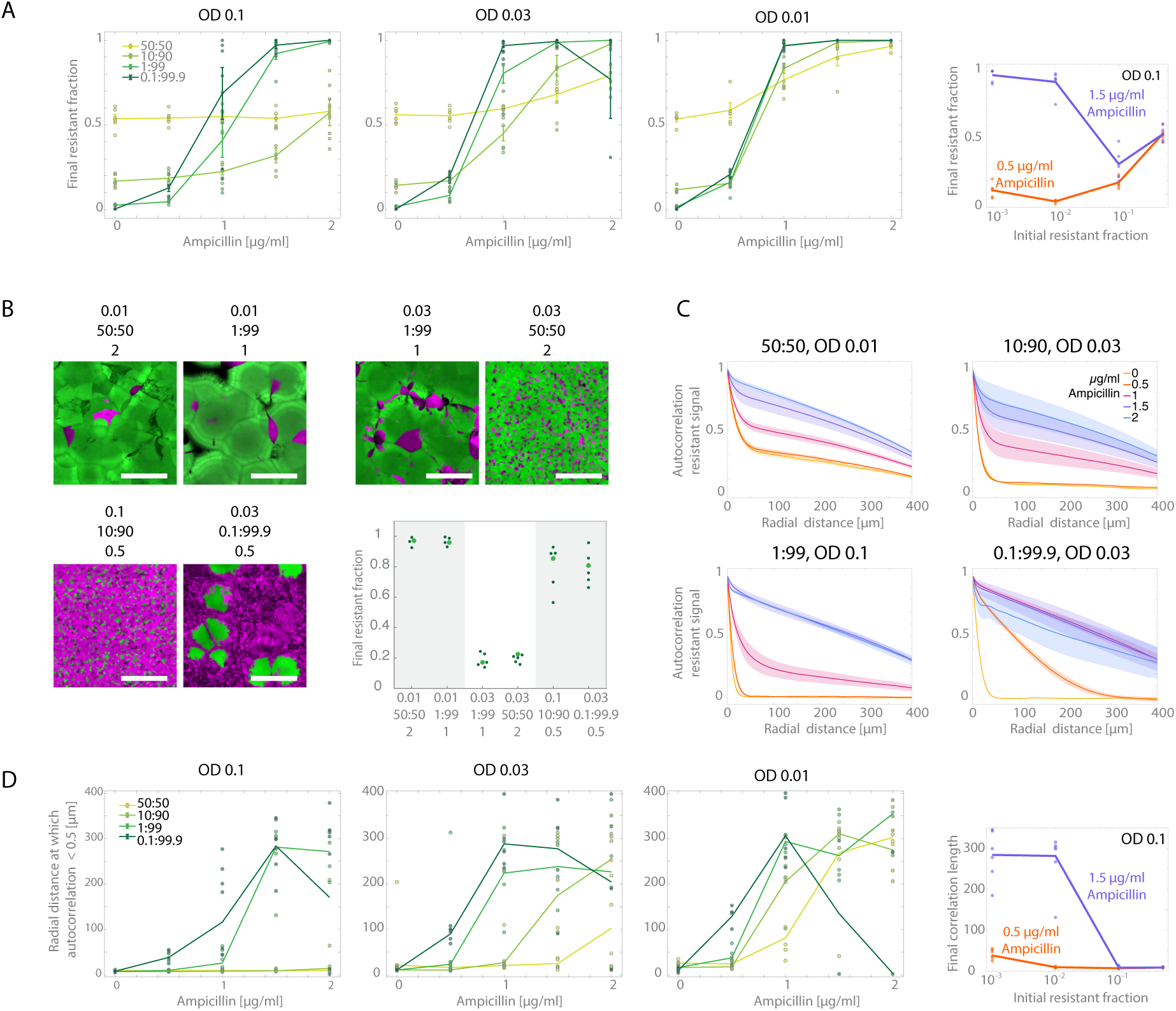
Homeland spatial organization changes composition and scale with initial conditions. (A) Final resistant fraction (segmented 2D resistant (“green”)-labeled area / (resistant+sensitive (“magenta”) area)) of a 1339.65 µm x 1339.65 µm image within the homeland, across different initial conditions. Mean and standard error, as well as individual replicate data, are shown. n= 3-6, see Figures S1-S3 for image data of all replicates. (B) Specific example pairs of patterns grown from different initial conditions, but ending at similar final fractions. Panel labels indicate initial density (OD), resistant:sensitive ratio, and ampicillin concentration from top to bottom. Scale bars 500 µm. Ampicillin-resistant bacteria in green, ampicillin-sensitive in magenta. Graph shows final resistant fractions across replicates for the shown conditions, with the larger, bright green data point representing the resistant fraction in the shown image. This is a subset of the microscopy data also shown in Figure 1 and the quantification in Figure 2A. (C) Radially averaged spatial autocorrelation functions (see Materials and Methods – Image Analysis) of segmented resistant/”green” signal for select initial conditions, used as a measure of pattern size with increasing ampicillin concentrations. Mean line and standard error as shaded area are shown. Whole dataset for all conditions is shown in Figure S6. (D) Distance at which radial correlation functions drop below 0.5 (“correlation length”) is used to describe pattern size (or scale) for all measured initial conditions. Mean curve and individual replicate data are shown.

In the presence of ampicillin, the final resistant fraction generally increased with ampicillin concentration, suggesting stronger selection for resistance and decreased cooperative protection of sensitive bacteria at higher concentrations. However, the degree of this increase was dependent on the initial resistant fraction and density, leading to a number of surprising results. At low ampicillin concentrations, increasing the initial fraction of resistant cells typically led to an increase in the final resistant fraction; for example, at OD= 0.1 and [AMP]= 0.5 µg/mL, increasing initial resistant fraction from 0.01 to 0.5 led to an increase in mean final resistant fraction from 0.04597 to 0.5403 (Figure 2A, left and right panel). Somewhat counterintuitively, we observed the opposite trend at higher ampicillin concentrations: seeding populations with increasingly resistant inocula led to *smaller* resistant fractions in the homeland. For example, at OD= 0.1 and [AMP]=1.5 µg/mL, the same increase in initial resistant fraction as above (from 0.01 to 0.5) led to a decrease in mean final resistant fraction from 0.9212 to 0.5388 (Figure 2A, left and right panel). In general, the relationship between initial resistant fraction and final resistant fraction was surprisingly complex and even non-monotonic at intermediate concentrations (Figure 2A).

In addition, we found that for sufficiently large initial resistant fractions, the final composition was essentially independent of ampicillin concentration (Figure 2A). For example, at a total density of OD=0.1, equal mixtures of sensitive and resistant cells gave rise to approximately 50:50 mixtures in the final colony, over the entire range of ampicillin concentrations tested (Figure 2A, left panel). The sensitivity to ampicillin concentration was partially restored, however, when the initial seeding was performed at lower densities (e.g. OD= 0.01, Figure 2A center right panel). At lower overall or resistant densities, we observed increasingly sharp transitions between initial ampicillin concentrations at which sensitive growth was protected, and slightly higher ampicillin concentrations at which protection was lost and resistant growth dominated. This effect is presumably caused by larger initial distances between bacteria in the colony, making protection less effective.

### Initial conditions modulate length scales of spatial organization

Although resistant fraction is an important metric, it does not capture all aspects of a spatially structured population. In our experiment, colonies with very different initial conditions at times reached similar final resistant fractions. However, colonies with similar final compositions often exhibited strikingly divergent pattern appearances (Figure 2B). Most obviously, the size of patches and degree of segregation of the two strains differed visibly, across an order of magnitude in some cases. To quantify trends in pattern size, we measured the radially averaged spatial autocorrelation of the resistant signal, which has a steeper slope when the resistant patch distribution is at a smaller length scale (and vice versa). Holding the initial resistant:sensitive ratio constant, the correlation scale tended to be larger with increasing ampicillin dosage and lower initial OD (Figures 2C, D and S6), in agreement with the visual impression that resistant spots tended to become bigger and take up a larger fraction of the area (Figure 1). In the absence of ampicillin, the scales of resistant patterns at the same initial density were similar, even when the initial ratios differed (Figure 2C), and when ampicillin was present, colonies with smaller initial resistant fractions consistently created larger resistant patches (Figure 2C, right panel). This was in contrast to the non-monotonic behavior of final composition with initial resistant fraction, and explains why the two measures don’t always correlate. Thus, the composition and scale of spatial patterns that form in the homeland under antibiotic exposure are shaped by non-trivial interactions between the initial properties of the colony and its environment.

### Cooperation acts at an interaction scale that is dynamic with resistant cluster growth

To understand how these trends in spatial organization emerge, we asked how ecological interactions between the two bacterial subpopulations scale across the colony. To exclude the possibility that cooperation is globally uniform across the agar plate, we showed that - for the same total initial cell numbers - sensitive bacteria were protected from antibiotic-induced death when intermixed with resistant bacteria inside a single droplet, but not when placed in a separate droplet on the same plate (Figure 3A). This suggests that protection does not extend across the entire plate.

**Figure 3:**
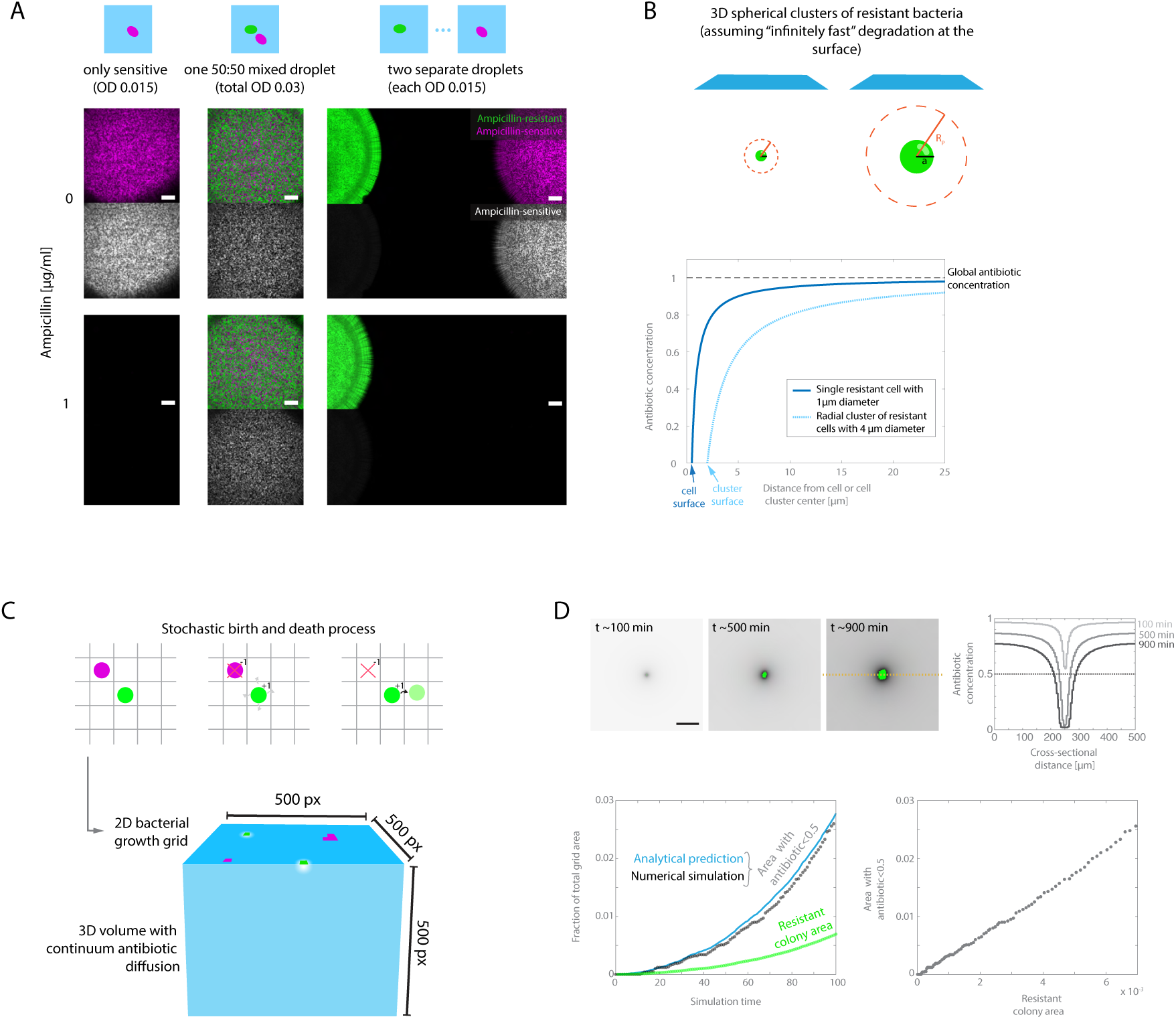
Antibiotic degradation and protection of sensitive bacteria occur at a dynamic length scale. (A) Protection of sensitive bacteria from antibiotic is distance-dependent and thus not a global effect. Microscopy images of either single sensitive (magenta) only colonies (left), single mixed resistant (green) and sensitive (magenta) colonies (center) or separate resistant and sensitive colonies (right) on agar with or without ampicillin. All samples contain the same density of sensitive bacteria. Scale bars = 500 µm. (B) Range of antibiotic gradients scales with resistant cluster size according to a steady state model. Illustration of mathematical prediction that spherical clusters of resistant cells can create larger equilibrium gradients of antibiotic than single resistant cells, assuming infinitely fast degradation and fixed global antibiotic at infinity, even if degradation only occurs at the location of resistant cells (no secretion and diffusion of enzymes). Graph shows quantitative analytical predictions of antibiotic profiles around a single resistant cell or a cluster of resistant cells, according to equations from ^57^. (C) Setup of the computational simulation used to numerically model spatial pattern formation in the homeland. Antibiotic-resistant (green) and -sensitive (magenta) bacteria are modeled as pixels on a two-dimensional grid (500x 500 pixels, ∼1µm or 1 cell diameter per pixel), and experience stochastic growth and death (sensitive only). Competition is modeled as restriction of growth to empty nearest neighbors; protection is modeled as a sensitive death rate that depends on the local antibiotic concentration. Antibiotic diffusion and degradation are modeled in three dimensions on a coarse-grained grid (500x500x500 µm^3^, 5x5x5 µm^3^ voxels) according to a forward time central space scheme. The cell grid corresponds to the top layer of the antibiotic volume and antibiotic is degraded at locations occupied by resistant bacteria. (D) The numerical simulation closely follows the analytical prediction for dynamic scaling of antibiotic depleted areas around resistant clusters. Top row: Timepoints of a growing antibiotic-resistant cluster from a single resistant cell (green) and antibiotic concentration (grayscale, showing the top layer of the antibiotic grid), with a global antibiotic concentration corresponding to 1µg/ml ampicillin, and corresponding cross-sectional antibiotic profiles (at the position indicated by the yellow line). Scale bar = 100 pixels = 100 µm. Bottom row: Increasing resistant cluster area and antibiotic depleted area (< 0.5 µg/ml) over time (left), and correlation between cluster area and antibiotic depleted area (right).

The steady-state protection radius offered by a single resistant colony is straightforward to calculate from a simple biophysical model. Specifically, if we assume that (fast) antibiotic degradation occurs on the colony surface, the distance-dependent antibiotic concentration surrounding a single resistant colony of radius a is given (for a 3D spherical colony) by 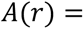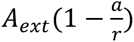^57^, where A_ext_ is the global initial antibiotic concentration. Therefore, the range of protection (i.e. the radius at which the antibiotic is reduced below a certain level, such as the MIC) is only a few micrometers for a single resistant cell, but linearly depends on the radius of the resistant cluster (Figure 3B) ^57^.

While this steady-state model may be sufficient for describing simple experiments involving a single resistant colony, growth in the homeland in multiple resistant patches causes potentially transient dynamics driven by multiple interacting antibiotic sinks. This suggests that protection length scales change dynamically and non-trivially throughout the experiment. We therefore need to go beyond steady state models with fixed interaction lengths to understand the growth and pattern formation process in the colony.

On the other hand, competitive interactions appear to be dominated by short-range effects. Within a colony, bacteria tended to fill the whole 2D homeland space and then maintain a stable pattern over several days (Figures 1 and S7), suggesting that competition is strong when bacterial growth fronts physically encountered another sub-colony. To understand whether long-range nutrient competition, which is an important interaction in other bacterial pattern formation systems ^72^, is a dominant factor here, we tested the colonies’ sensitivity to variation in overall nutrient concentration. Reducing nutrient concentration slowed overall growth but did not seem to affect pattern formation, whereas increasing nutrient concentration three-fold appeared to moderately increase the sensitive fraction at 1µg/ml Ampicillin (Figure S8). This could be explained by higher nutrient concentrations reducing competition and thus increasing the growth rate of all cells, including resistant clusters, relative to the death rate of sensitive cells. Thus, a large protective effect might be achieved more quickly in the growth process.

However, since pattern formation in 2D did not appear to be very sensitive to nutrient concentrations, we focused on the interplay between competition for space and cooperative antibiotic resistance. Based on our observations, we hypothesized that the spatial organization in this system may be driven by an interplay between local competitive and dynamically scaled protective interactions between the antibiotic-resistant and -sensitive subpopulations.

### A biophysical model with short-range competition and dynamic-range cooperation recapitulates observed spatial patterns

To test whether emergent dynamics from two ecological interactions interplaying across space and time was sufficient to create the types of spatial arrangements we observed in our experiments, we built a numerical model to simulate the growth process (Figure 3C). We modeled a 500-by-500 µm^2^ subregion of the bacterial colony homeland (∼4,000 µm diameter) as a 2D grid with periodic boundaries, with one grid point corresponding to one cell or about 1 µm^2^. Antibiotic-resistant and -sensitive “bacteria” on the grid performed stochastic birth and death (sensitive only) processes similar to an Eden-style model ^73^, whose probabilities depended on the available space and the local antibiotic concentration, in time steps corresponding to about ∼10 min (∼1/2 division cycle) (See Materials and Methods for a more detailed model description). Spatially resolved antibiotic concentration was modeled by continuum diffusion in a 3D volume (500x500x500 µm^3^) with a coarse-grained grid (5 µm edge voxels), one surface of which corresponded to the colony grid. Antibiotic was fully degraded at locations corresponding to resistant cell positions, and then diffusion was modeled according to a Forward Time Central Space scheme ^74^. After 3D diffusion and degradation steps adding up to ∼10 minutes, the final antibiotic landscape was used to compute the next growth and death step on the 2D colony grid. Thus, our model contained two primary ingredients: degradation of antibiotic by resistant cells (giving rise to time-dependent protection zones as resistant cells reproduce) and local (nearest neighbor) competition for space. We used literature values to approximate baseline bacterial growth rates and ampicillin degradation and diffusion speeds, and fit the sensitive death rate to separate dose-response curve experiments of sensitive-only populations (See Materials and Methods, Figure S9A), rendering the model without free parameters.

We first confirmed that this model replicated a similar scaling relationship between resistant cluster size and antibiotic degradation as was predicted analytically for the steady state (Figure 3D). As the simulated patch grew, the area in which the antibiotic was depleted grew as well; the protection zone (i.e. the area with antibiotic concentration below half the global concentration) closely matched the analytical prediction, and appeared to have a linear relationship with resistant cluster area (slope approximately 4 for an initial global concentration of 1 µg/ml ampicillin).

We then sought to replicate the experimentally observed pattern formation of mixed antibiotic resistant and sensitive bacteria. Despite the simplicity of the model, it qualitatively reproduced multiple features of the experiment, including spatial patterns that were strikingly similar to the experimental microscopy images (Figures 4A and S10-S12). This qualitative agreement held true across a wide range of tested conditions, including different ampicillin concentrations and initial colony densities and resistant fractions, and the model even quantitatively captured population compositions (Figure 4B) and correlation lengths (resistant cell cluster sizes; Figures 4C and S13) over a wide range of initial conditions. We do note some quantitative disagreement—for example, in some cases the model overestimated sensitive cell survival—although even in these cases, the model often matched closely with at least one of the experimental replicates (Figures S10-S12). We stress that protective interactions were required to achieve patterns reminiscent of the experimental data; a simulation without active degradation of antibiotic, in which interactions between the subpopulations are purely competitive, did not reproduce the experimentally observed patterns (Figure S9B). Overall, these results suggest that our experiments are dominated by local competition and dynamically-scaled cooperation between antibiotic-resistant and sensitive bacteria.

**Figure 4:**
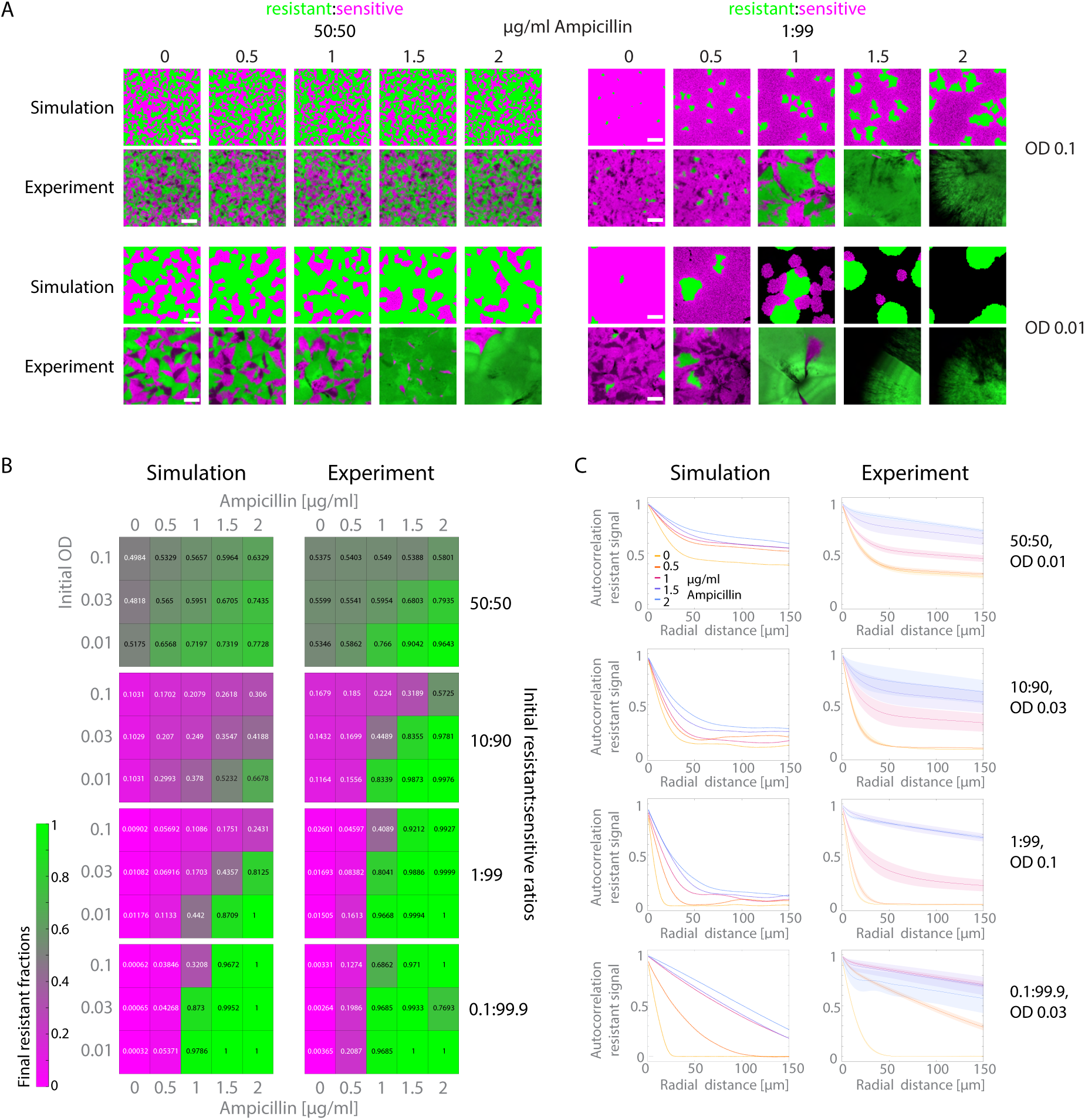
A simple cellular automaton model with cooperation and competition can reproduce many features of spatial organization of resistant and sensitive subpopulations. (A) Simulation produces spatial arrangements reminiscent of those observed in experiments. Simulation results for cell grids and corresponding experimental images for select conditions. Scale bars = 100 µm. Antibiotic-resistant cells in green and antibiotic sensitive cells in magenta, respectively, no-occupied areas of simulation in black. Full dataset comparisons in Figures S10-12. Experimental images shown are 500x500 µm^2^ representative subregions of data shown in Figure 1. (B) Simulation trends in colony composition are similar to experiment quantifications. Heat maps showing the resistant fraction encoded as a color map (more green = higher resistant fraction, more magenta = lower resistant fraction) across all simulated and experimentally tested conditions. Number in each square denotes the resistant fraction (n = 1 for simulations, n = 3-6 for experiments, mean is shown – same experimental quantification as shown in Figure 2A). (C) Comparison of radially averaged autocorrelation functions of resistant/green signal for simulations and experiments across select initial conditions. This metric compares the spatial scale of patterns, with more coarse patterns (larger patches) maintaining higher correlations at longer distances. Mean line and standard error as shaded area are shown for experiments, same experimental quantification as shown in Figure 2C. Full dataset in Figures S13.

### Spatial interplay of protection and competition drives complex spatial growth dynamics of resistant and sensitive subpopulations

We next took advantage of the simulation data to get a better intuition for how the ecological dynamics of resistant and sensitive bacteria result in the complex and diverse final spatial arrangements that we had observed across different conditions. One prominent aspect of these spatial patterns were drastic transitions between majority-sensitive and majority-resistant patterns that occurred with only small changes in external antibiotic concentration. To understand the relative dynamics of resistant growth, antibiotic degradation and sensitive growth and death, we first quantified the spatial area covered by each subpopulation and the total area of “protection zones” (antibiotic below 0.5 µg/ml) over time (Figures 5A, S14). The simulations suggested that even a relatively small area of resistant cell coverage was capable of driving the antibiotic concentration below the threshold concentration across the entire simulated surface. The antibiotic-cleared area clearly diverged from proportional increase with resistant growth (expected from steady-state models) in those cases (Figure 5A), emphasizing the dynamic nature of the length scale of protective interactions. Specifically for colonies seeded with high resistant fractions, transition to clearance of the entire surface occurred rapidly in the early stages of growth, which suggests why, both in simulation and experiments, patterns similar to the antibiotic-free conditions emerged. When the initial resistant number was decreased—either with lower fraction or total density—and at higher ampicillin concentrations – which are expected to cause steeper antibiotic gradient around resistant sinks even when degradation is infinitely fast– , this transition occurred later in the growth process.

**Figure 5:**
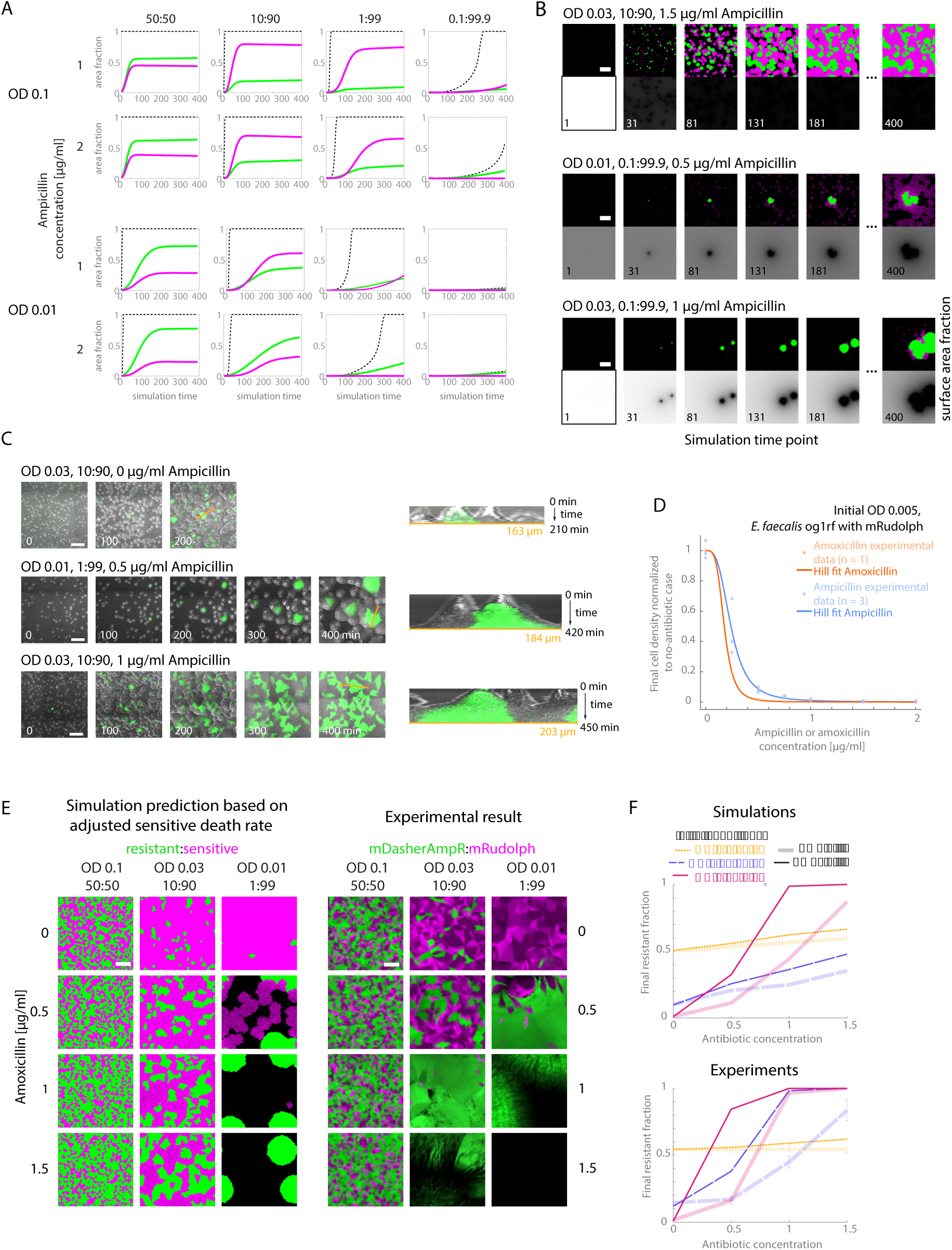
Simulations elucidate spatiotemporal population dynamics and predict spatial organization under novel conditions. (A) Simulations suggest complex and varied global population dynamics during growth of the colony homeland. Simulated resistant (green)- and sensitive (magenta)-covered, and antibiotic depleted (< 0.5µg/ml, black dashed line) area as a fraction of total grid area over simulation time, for representative initial conditions. Simulation dynamics for all conditions can be found in Figure S14. (B) Examples of simulated spatial dynamics of the two interacting populations across growth timepoints, from different initial conditions. Scale bars are 100 µm. Top row images for each condition show resistant (green) and sensitive (magenta) cells and bottom row images show the top layer of the three-dimensional antibiotic grid, with antibiotic concentration in gray-scale (white = 1 µg/ml or above, black = 0 µg/ml). Final images correspond to those shown in Figures 3, S10-12. (C) Experimental data showing snapshots of the spatial growth process over time. Transmitted light channel depicts the outlines of cell clusters in grayscale, ampicillin-resistant clusters are marked in green. Left: individual timepoints (labeled in minutes after the start of imaging). Scale bars = 100 µm. Right: Kymographs showing the corresponding colonies over time at the cross-sections indicated by orange lines on the left. Different growth rates of resistant and sensitive colonies, as well as competition at boundaries are observed. (D) Comparison of dose-response curves of the sensitive *E. faecalis* strain to ampicillin (blue) or amoxicillin (orange), a different antibiotic in the class of β-lactam antibiotics, while growing in a colony on agar, indicating higher sensitivity to amoxicillin. Experimental measurements (n = 3 for ampicillin, n = 1 for amoxicillin) and corresponding best-fit hill-like curves are shown. (E) Left: Simulation predictions for spatial organization using an adjusted sensitive death rate to match the amoxicillin dose response curve across different initial conditions. Right: Experimentally observed spatial arrangements using amoxicillin. Resistant bacteria in green, sensitive bacteria in magenta. Scale bars = 100 µm. (F) Quantification of shifts in resistant fractions for the same initial ampicillin vs amoxicillin concentrations, in simulations and experiments (mean and, if applicable, standard error are shown; n= 3-6 for ampicillin experiments, n = 1 for amoxicillin experimental and all simulations). Ampicillin quantifications are the same as shown in Figures 2A and 4B.

These trends coincided with an early decline in sensitive cells that sometimes led to extinction of the sensitive subpopulation. This interplay also illustrated why resistant patches tended to grow to larger sizes at higher antibiotic concentrations or lower resistant fractions (Figure 2), as sensitive patches are more likely to die out in those cases before experiencing sufficiently reduced local antibiotic concentration and thus do not provide much competition to resistant growth.

While these global dynamics of antibiotic provided a relevant predictor for bulk population composition, we found that early local dynamics of sensitive cell growth, driven by local antibiotic depletion around small resistant clusters, were also important to understand spatial pattern formation. The resulting local competition between sensitive and resistant cells drove spatial pattern formation, leading to rough, branched or undulating interfaces reminiscent of what we observed in experimental colonies (Figure 5B, Supplemental Movies S1-6). Specifically, the simulations suggested that resistant growth dominates in early stages of growth, but once resistant clusters reach sufficient sizes, they afford protection to neighboring sensitive bacteria, which then compete with resistant cells for space. This local competition caused the surface of the resistant cluster to branch out to grow into areas not yet occupied, resulting in fractal-like interfaces. Eventually, sensitive growth was often able to slow down and overtake resistant growth, despite the absence of an explicit fitness cost. However, at sufficiently high global antibiotic concentrations and low resistant numbers, only those sensitive clusters initially seeded close to a resistant clusters could even survive, leading to small sensitive enclaves within or adjacent to the resistant clusters, while the global sensitive subpopulation largely went extinct.

To investigate these predicted dynamics experimentally, we acquired time lapse movies of parts of the growth process for select experimental conditions (Figure 5C). These movies confirmed that, in the absence of ampicillin, resistant and sensitive subpopulations grew at similar rates and largely stopped growing in 2D at boundaries with other clusters due to local competition (Figure 5C, top panel). When ampicillin was present (Figure 5C, middle and bottom panels), resistant clusters grew fast at early stages followed by growth of neighboring sensitive clusters that eventually impeded resistant growth at interfaces. As resistant clusters encountered those interfaces, we also observed them growing into remaining open areas, causing branching by a mechanism consistent with our model predictions.

In summary, our time-resolved simulation and experimental data provided insights into how dynamic interplay of local competition and expanding protective interactions during the growth process could result in a wide variety of spatially organized colony homelands at different compositions and scales, with sometimes drastic transitions between types of spatial arrangements.

### Simulation can predict spatial population organization changes under novel conditions

We next investigated whether our model could be extended to predict spatial organization occurring under novel, different conditions. We first confirmed that the model was able to predict patterns formed at an intermediate ampicillin concentration (0.75 µg/ml) similarly well to the initially tested ampicillin concentrations (Figure S15). We then sought to predict the effect of sulbactam, a β-lactamase inhibitor that is often applied together with ampicillin to suppress resistance during clinical treatment ^75,76^. To mimic β-lactamase inhibition, we reduced the frequency of antibiotic degradation in our model (Fig S15). We found that reducing degradation frequency by a factor of 5 or 10 (see Materials and Methods for details) produced remarkably similar results to experiments with sulbactam concentrations of 1 and 2 µg/ml respectively, across different ampicillin concentrations and initial colony compositions. Because our model didn’t take into account resistant cell death, it was not able to reproduce the increased ampicillin sensitivity of resistant cells that is observed with sulbactam.

We then asked whether this model could also apply to the effects of a different, related antibiotic, the β-lactam amoxicillin (Figure 5D and E). As before, we estimated the sensitive death rate parameter using a separate dose-response experiment in amoxicillin, which showed a slightly steeper dose response curve than ampicillin (Figure 5D). The model and experiments both exhibited a notable change (with antibiotic concentration) in the composition of simulated patterns from ampicillin to amoxicillin, while qualitative features of the patterns stayed largely the same (Figure 5E, F). As before, we observed some quantitative discrepancies between the model and the experiments, but relative composition shifts from ampicillin to amoxicillin, in terms of both direction and magnitude, were captured remarkably well. Thus, our model has potential utility for other systems driven by antibiotic-mediated killing and antibiotic degradation, requiring only independent estimation of two effective parameters: sensitive cell death rate and antibiotic degradation rate.

### Homeland dynamics affect consequent range expansion behaviors

Range expansions are popular model systems for exploring ecological dynamics ^35,52,68,77,78^. Most studies of range expansions focus on growth outside of the homeland, neglecting potential effects that could arise from spatial dynamics in the homeland itself. To explore some of the functional consequences of spatial population dynamics in the homeland, we asked how homeland spatial organization may influence the subsequent range-expansion dynamics at the edge of the colony. Our previous experiments (Figures 1 and 2) demonstrated that the composition and distribution of resistant and sensitive subpopulations in the homeland could change drastically during growth under antibiotic exposure. We hypothesized that these homeland differences could impact range expansion dynamics by 1) modulating the composition and segregation of the homeland population, which serves as the starting point for range expansion, and 2) sustaining long-range protective interactions between the homeland and the expanding fronts of the range expansion.

Indeed, especially at high antibiotic concentration and low initial densities, our experiments had showed that population composition in the homeland can vary substantially, even in populations seeded with similar initial resistant fractions. These different homeland compositions appeared to also be associated with different segregation patterns in the range expansion region (Figure 6A, top and middle panels). Conversely, colonies with very different initial resistant fractions could produce similar homeland patterns and subsequently similar range expansion patterns (Figure 6A, bottom panel).

**Figure 6:**
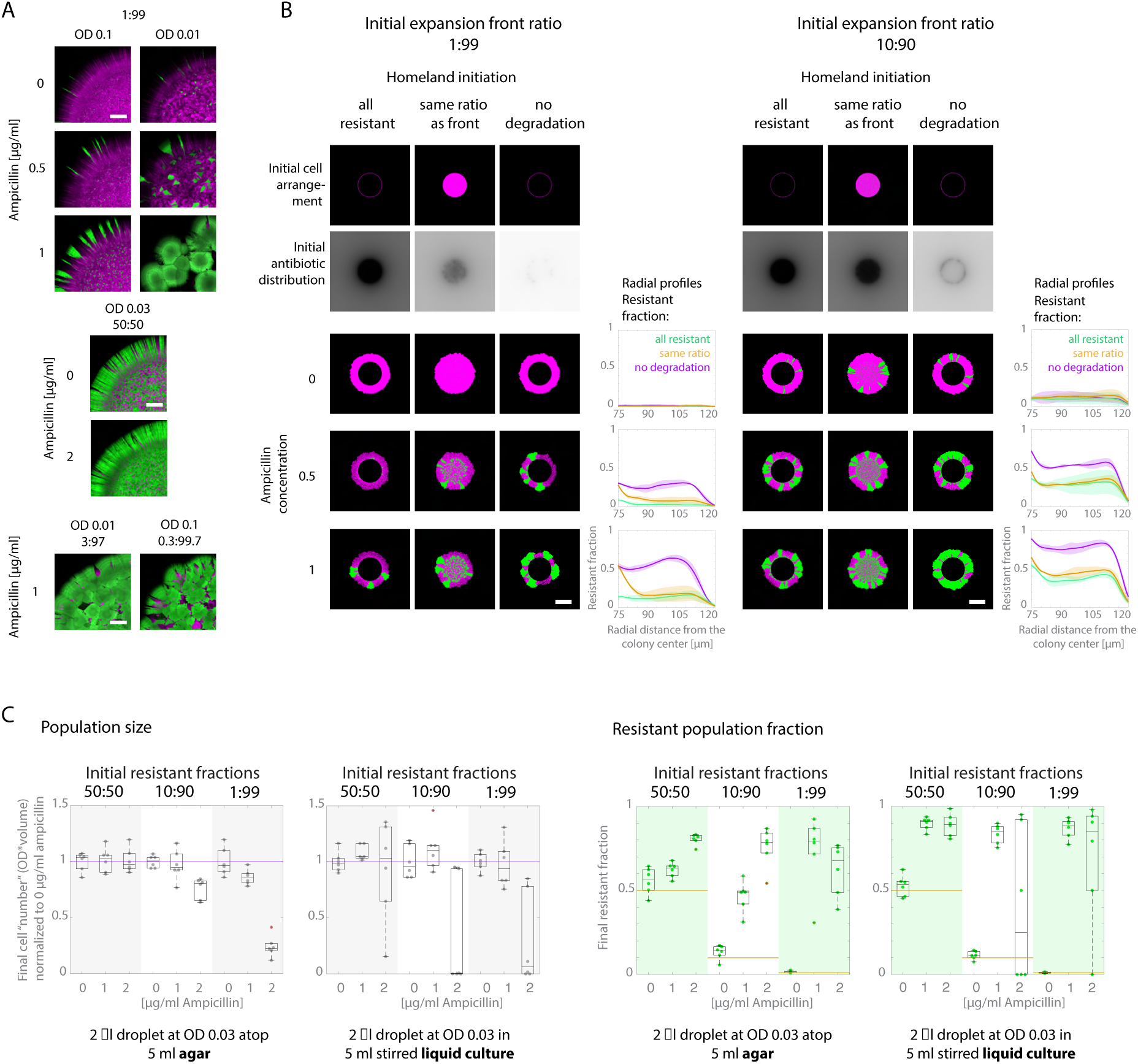
Spatiotemporal population dynamics of the homeland have functional consequences for the bacterial community. (A) Homeland population dynamics influence the initiation of the range expansion front. Confocal image maximum intensity projections showing the homeland (colony center is towards the bottom right of the image) and range-expanding region (towards the top left) after ∼18h of growth, showing about ¼ of the colony, across different initial conditions as labeled. Scale bars = 500 µm. Resistant bacteria in green, sensitive bacteria in magenta. (B) Simulations showing how degradation by resistant cells in the homeland could affect range expansion dynamics for two different initial resistant ratios at the expansion front and three different homeland compositions: all resistant/degrading, same ratio as the expansion front or non-degrading. In the all-resistant and no degradation cases, homeland cells do not contribute by growing or dying, only to degradation of antibiotic if applicable (non-growing homeland cells are not shown explicitly in images; they do block inward growth of the range expanding front). Initial cell arrangement and final range expansion patterns of resistant (green) and sensitive (magenta) bacteria for different global ampicillin concentrations are shown, as well as the antibiotic distribution at the initial time step (n = 3 simulations for each conditions, all individual images can be found in Figure S16). Scale bars = 100 µm. Radially averaged resistant fractions (of total line area at a certain radius, including empty regions) along the range expansion area are also shown for each homeland condition, showing that composition of the range expanding region is affected by homeland-mediated degradation of antibiotic. Mean lines and standard deviation as shaded area - after 5 µm moving average smoothing was applied to each replicate - are shown. (C) Comparison of normalized *E. faecalis* population sizes and resistant fractions in agar colonies versus well-mixed liquid cultures. Left and left center panels show relative final population sizes after ∼18 h of growth at 37°C under ampicillin exposure (normalized to respective 0 µg/ml ampicillin condition mean) for populations grown on agar (left) versus in well-mixed liquid culture (left center), at different initial resistant:sensitive ratios (indicated by shaded areas), all with an initial OD of 0.03. Purple line indicates the mean final population size of the 0 µg/ml ampicillin condition, for each initial population composition. Right center and right panels show final resistant fractions (measured by green fluorescence for a sample of each culture under the microscope) for the same experiments. Yellow lines serve as visual markers of initial resistant fractions for each experiment. Box plots with median and quartile lines and whiskers indicating data spread (apart from outliers) are shown, as well as individual data points (n = 6 for each condition). Outliers are defined as values more than 1.5 interquartile ranges from the edges of the box and shown as red ‘+’. Non-normalized population sizes are shown in Figure S17.

To investigate the impact of long-range interactions between the homeland and the range expansion region separately from any composition changes in the homeland, we simulated a series of three matched range expansion fronts (starting from the same initial composition) radiating from non-proliferating homeland populations with different antibiotic-degrading properties (Figures 6B and S16). In one case, the cells in the homeland acted like resistant (they degrade antibiotic) but non-growing cells. In the second case, cells in the homeland acted like non-growing/dying sensitive cells (they did not degrade antibiotic). In the third case, cells in the homeland contained a mixed population whose composition (and growth/death rules) were identical to the initial composition of the range expansion population. We observed that range expansion simulations with the same initial front composition diverged drastically depending on whether the homeland was filled with non-degraders (sensitive cells), all degrading cells (resistant cells), or with a population of the same composition as the expansion front. When protection stemming from the homeland was high (e.g. “all resistant” case), sensitive bacteria at the expansion front experienced lower local antibiotic concentrations and were thus better at competing with resistant sectors. Thus, the resistant fraction throughout the range expanding region was decreased by the influence of the homeland. This effect was robust across two different initial resistant fractions and amplified with higher global antibiotic concentration. The resulting altered population composition of the expansion front likely further affects range expansion, even at distances outside of the range protected by the homeland, creating a sustained effect. Overall, our results suggested that ecological interactions both within the homeland and between the homeland and the expanding population may drive outcomes in range expansion experiments.

### Spatially-organized colonies have different community-scale outcomes than comparable well-mixed cultures

Our results suggested that spatial interactions between sensitive and resistant cells can give rise to a wide range of population dynamics in the homeland. How do these compare to the dynamics of similar populations in well-mixed liquid cultures? While ecological models frequently neglect spatial structure for simplicity, it is unclear to what extent it was an important factor in driving this system’s response to antibiotic. To answer this question, we compared population size and composition of matched populations grown either 1) on agar or 2) in well-mixed cultures (Figures 6C and S17). We found that, at low initial resistant fractions and high antibiotic concentrations, liquid cultures displayed bimodal outcomes, either reaching population sizes similar to the no-antibiotic case, or “crashing” to extinction. These findings are consistent with previous work in continuous flow bioreactors, where populations exhibit bi- stability between survival and extinction ^36,79^. In contrast, agar colonies consistently grew to an intermediate size, somewhat below their antibiotic-free population size, suggesting that the two culture types exhibit fundamentally different population responses to ampicillin exposure. One possible explanation is that on agar, resistant cells can clear small, local areas from antibiotic, which may be sufficient for even just subsections of the colony with high local resistant density to survive and grow; by contrast, in liquid culture, the entire global antibiotic concentration needs to be reduced in order to improve survival of the dispersed well-mixed population, which may make it sensitive to small variations in resistant cell number or growth rate.

We also found that population composition was impacted by spatial structure. While the resistant fraction tended to increase with ampicillin concentration for all conditions, the increase in resistant fraction was typically larger in liquid culture than on agar, suggesting a stronger selection for resistance in well-mixed communities, consistent with recent findings in *E. coli* biofilms ^32^. This can be explained again by the fact that on agar, sensitive cells can be protected locally, even before any significant reduction of global ampicillin concentrations has occurred, and can then also compete with resistant growth. In fact, in agar colonies, homeland spatial patterns and compositions were not affected by changing the total agar volume, suggesting that those colonies grow independently of the global ampicillin supply (Figure S18).

In summary, we observed differences between the outcomes of spatially structured population growth, in which local and spatially restricted ecological interactions are more dominant, and well-mixed population growth, in which global interactions are expected to be more important. This underscores the importance of studying collective antibiotic resistance and ecology of microbial communities in spatially structured systems, which make up a majority of bacterial life ^3^.

## Discussion

We have described how spatiotemporal population dynamics of ampicillin-sensitive and - resistant *E. faecalis* subpopulations result in emergent spatial organization in the homeland of colonies. Varying the initial colony composition and antibiotic concentration produced a surprisingly broad range of spatial arrangements in the homeland of the colonies, with often rapid transitions between only slightly differing conditions. Dissecting how the ecological interactions between subpopulations act in a spatially heterogeneous manner allowed us to mechanistically understand and predict the resulting composition and spatial structure of these populations under antibiotic exposure. We also explored ways in which these spatially scaled interactions have consequences for population behavior relative to well-mixed systems and for subsequent range expansion dynamics. These findings emphasize the importance of spatial structure and its effects on population dynamics for antibiotic resistance.

Our experimental results could be largely explained with a simple biophysical model that incorporates local competition and (antibiotic) diffusion-driven protective interactions between the populations. Importantly, our results highlight that the range of protection from resistant subpopulations is not at a set length scale, but increases dynamically as the clusters grow, leading to non-trivial coupling between the growth process, the population composition, and the local antibiotic concentration. As a result, phenomenological models that assign a fixed length scale to cooperation—for example, models that include a local Allee effect ^52^ restrict interactions to nearest neighbors on a fixed graph structure ^80,81^ — may be insufficient to describe our results. This dynamic scaling helps explain why the spatial patterns of resistant and sensitive patches resulting from these growth and death processes have such a wide range of characteristic length scales.

It is important to keep in mind several limitations of our work. First, our model is, of course, a dramatic oversimplification of the underlying biology driving biofilm formation. For example, we assumed competition reduces growth by preventing cell division whenever neighboring space is unavailable, yet colonies are expected to have active growth zones of at least a few cell diameters and may physically push away neighbors within those zones ^82–84^. However, our data does show that growth in the 2D plane of the agar plate is dramatically and suddenly reduced when another cluster is encountered (Figure 5C), suggesting that local competition is a dominant aspect of this system. This restriction contributed to the model needing to run for more time steps to achieve comparable area coverage to the experiments. Further, we neglected a number of mechanisms known to contribute to antibiotic responses, such as persister cells, tolerance, or the evolution of new resistance mechanisms ^85–88^ (in part due to the short time scale of our experiments). We also did not take into account effects due to cell shape and cell motility, since *E. faecalis* is spherical and immotile, but which are likely relevant for some microbial species. For example, rod-shaped bacteria such as *E. coli* or *Bacillus subtilis* form winding fractal patterns driven by their elongated cell-scale shapes ^64,65^; similarly, antibiotic-induced filamentation has been shown to impact the ecological interactions in microbial communities ^37^. Additional, antibiotic-independent ecological interactions, such as cross-feeding mutualism, can also drastically influence spatial organization of resistant and sensitive bacteria ^89^ in ways that feed back into the dynamics of antibiotic protection ^90^. Previous work in other microbes (and without antibiotics) has shown that different spatial patterns of interacting (sub-)populations can emerge from various types of intercellular interactions ^91^, initial partitioning of the seed population ^55,92^, or intercellular adhesion ^93^. In addition, recent work highlights time-dependent scales of cooperation in motile bacteria, reminiscent of the variable-range cooperation we observe in non-motile *E. faecalis* ^94^. The statistical inference model introduced there may offer an alternative approach to investigate these dynamic length scales in our system. Finally, we considered bacterial growth as a 2D process, but recent advances highlight that interesting dynamics can also occur in the vertical direction of growing bacterial colonies in other species ^95,96^.

We applied our model to compare two similar antibiotics (ampicillin and amoxicillin) and also to combination treatments involving an antibiotic and enzyme inhibitor. Nevertheless, it is not certain that the model will translate to other systems, even other cooperative antibiotic resistance mechanisms. In addition, our simulation assumes that the degrading enzyme stays bound to the resistant bacteria and doesn’t diffuse into the rest of the colony. This appears to be true for *E. faecalis* ^62,63^, and chloramphenicol acyltransferase in *Streptococcus pneumoniae* also conveys protection by intracellular degradation activity ^24^. However, β-lactamases in other species are thought to be secreted ^97,98^, which could change the dynamics of protection.

Nevertheless, the processes in our model are not necessarily mechanism- or species-specific, requiring only estimates of death rates and antibiotic degradation rates, and thus this model could be applicable more broadly.

Our results are complementary to other studies investigating pattern formation and ecological interactions—including those mediated by public goods—in spatially heterogeneous communities ^4,47^. In our case, the “public good” is the lowered antibiotic concentration around resistant degraders, but there is also a significant body of work investigating other types of public goods in microbiology—for example, cooperators that secrete valuable metabolites that are consumed by non-producing cheaters ^99^. While conventional wisdom suggests that spatial structure aids cooperators by keeping their public goods restricted to kin ^47,50–54^, we observed that the antibiotic-resistant cooperator strain retains a small population fraction in spatial relative to well-mixed colonies. This is in line with other recent work suggesting more nuanced effects of spatial structure on cooperation ^100^.

Finally, although we only focused on ecology of two subpopulations with fixed genetic identities, our findings could have interesting implications for other aspects of eco-evolutionary dynamics, especially at longer time scales. Since de novo antibiotic resistance may be enabled by delayed death in protection zones ^72,100^, or can even evolve faster on antibiotic gradients ^101,102^, it is conceivable that the heterogeneous antibiotic landscape created by resistant clusters promotes resistance evolution in the sensitive subpopulation. And while the plasmids used here do not appear to transfer between subpopulations, horizontal gene transfer (HGT) can play an important role in the ecology of antibiotic resistance generally ^34,103^. The different interfaces between subpopulations generated in the spatial organization processes we describe here could affect HGT for competent plasmids or strains, thus potentially creating interesting feedback dynamics.

## Materials and Methods

### Bacterial strains and plasmids

*E. faecalis* og1rf strains transformed with pBSU101 plasmids ^104^ containing fluorescent labels and a spectinomycin resistance gene, some of which contained a blaZ ampicillin resistance gene encoding a β-lactamase from a clinical CH19 strain ^28^, were used for these experiments, as previously described ^57,69^:

| Strain name(s) | Fluorophore & Excitation wavelength | Antibiotic resistance genes |
| --- | --- | --- |
| "mRudolph"/"sensitive" | mRudolph (ProteinPaintbox)-optimized, 555 nm | spc (Spectinomycin resistance gene) |
| "mDasherAmpR"/"resistant" | mDasher (ProteinPaintbox), 488 nm | blaZ (Ampicillin resistance) and spc |
| "mRudolphAmpR" | mRudolph (ProteinPaintbox) - optimized, 555 nm | blaZ and spc |
| "mDasher" | mDasher (ProteinPaintbox), 488 nm | spc |

### Growing agar colonies under antibiotic exposure

Single colonies of relevant strains were inoculated into ∼3 ml Brain Heart Infusion (BHI) media (Remel) with 120 µg/ml spectinomycin sulfate (Thermo Fisher Scientific, 430.43 g/mol, standard concentration for all experiments unless otherwise noted) to select for pBSU101 plasmids over night at 37°C. Overnight cultures were then diluted 1 in 100 in fresh BHI + spectinomycin and grown to exponential phase (∼2.5-3 hours, OD600 between 0.1 and 1). 35 mm diameter petri dishes containing 5 ml of 15 g/L agar were freshly prepared the same day, mixing in 120 µg/ml spectinomycin and 0- 2 µg/ml of ampicillin sodium salt (Fisher BioReagents, 371.39 g/mol) from frozen stock solutions after the agar had cooled below 55°C in a water bath. Note that antibiotic concentrations are reported throughout this paper with respect to the weight of the respective salt, not the “pure” antibiotic molecule. Optical density (OD) measurements were performed on exponential phase cultures, and then mixtures of ampicillin-resistant and -sensitive strains at specified total densities and fractions were calculated and prepared (with densities approximated as OD, assuming linear relationship): Volume of culture in mixture = Goal volume total x Fraction(strain) x OD(goal)/ OD(measured). 2 µl of a mixed culture were pipetted onto the surface of an agar plate (one per plate unless otherwise stated) and let dry inside a laminar flow hood. All plates were then placed upside down at 37°C (at high humidity) for about 18 hours to allow colonies to grow and fill the homeland (Figure 1A).

Colonies grown without spectinomycin in the agar formed similar patterns, apart from increased frequency of non-fluorescent patches as the plasmids were not selected for to the same extent, suggesting that there was no obvious drug interaction between ampicillin and spectinomycin confounding our observations (Figure S19). Although there were small differences in the exact patterns when comparing different strains labeled with swapped fluorophores, likely due to small variations in growth rate and ampicillin resistance, the overall trends and pattern types were consistent (Figure S20).

To test the scale of cooperation, we also performed an experiment where two droplets were seeded several millimeter apart and the let grow overnight (Figure 3A). We additionally tested the effect of agar volume and height (and thus total ampicillin) by growing colonies on a 35mm agar dish with 3ml agar and 10cm dishes with 20ml agar (Figure S18). To test nutrient dependency, we prepared agar with different BHI concentrations between 0.1x and 3x the recommended concentration (Figure S8). To test pattern formation under amoxicillin (Figure 5), a low concentration solution of amoxicillin trihydrate (RPI Research Products, 419.5 g/mol) in water was prepared the day of the experiment and then used immediately to prepare plates at final amoxicillin concentrations of 0 to 2 µg/ml.

### Confocal imaging

To observe the spatial arrangement of bacteria inside an agar colony, a custom 3D printed spacer was placed on top of the agar, keeping a 18x18-1.5 cover glass (Fisherbrand) about 0.5 mm away from the agar surface where it did not perturb the colony, but not far enough to bring the colony beyond the focal length of the objective (Design file available at 10.5281/zenodo.14226311). Colony overview images were acquired on the Zeiss LSM 700 inverted confocal microscope with a Plan Apochromat 20x/0.8 M27 objective lens, with the pin hole at maximum opening (460.5672), using the 488 nm and 555 nm lasers to excite mDasher and mRudolph fluorophores, and detector ranges of 0-550 nm and 560-1000 nm, respectively. Either the full area of the homeland, or at least a 2.9 mm x 2.9 mm area within the homeland, was imaged using tile scans, at x and y pixel sizes of 0.8931 µm. A stack of 3 and 12 z-slices (13.358 µm each) was typically sufficient to capture the colony surface across the xy extent of the image.

For images displaying more details at the scale of individual cells, a smaller pinhole size (32.549) and smaller xy pixel (0.3126 µm) and z-step (0.9833 µm) were used. Such data was used to approximate the height of the biofilm and to show that the boundaries of the spatial patterns were largely constant across the depth of the colony (Figure S4). Because our imaging method did not disrupt the colonies, we were able to put plates back into the incubator and examine how patterns changed several days later (Figure S7). For 10 ml agar dishes, a ∼3-4cm diameter area around colony had to be cut out after overnight growth and flipped onto a 3D printed disk that had a hole fitting the imaging window.

### Image analysis

Tile scans were assembled using the “Stitching” plugin ^105^ in FIJI/ImageJ ^106^ with positions from metadata, linear blending and not computing overlap. A maximum intensity z-projection was applied across all z-positions of the assembled files to display the 2D pattern across the colony surface. A 1339.65 µm x 1339.65 µm (1500 pixels x 1500 pixels) representative area of the homeland was selected for image processing, since the patterns were largely uniform across the homeland, and this allowed for faster image analysis. The “maximum” of each channel was automatically adjusted to an optimal brightness, using the Brightness/Contrast > Auto function in FIJI/ImageJ, and then green (mDasher, 488 nm) and magenta (mRudolph, 555nm) channels were combined to RGB images and exported. Because the magenta channel had some background signal, the maximum adjustment was limited to stay above 30 (out of 255). Since we weren’t measuring relative intensities and there was no real overlap between the populations, this image processing pipeline was optimized to achieve accurate segmentation rather than maintaining consistent brightness information.

These images were segmented then into green (“resistant”), magenta (“sensitive”) and empty areas (no fluorescence or no bacteria) using a machine learning interface, ilastik ^71^, that was trained on four representative images of colonies with different compositions and verified to closely match visual interpretation of the images. The segmentation results were used to calculate resistant fractions (green/(green+magenta)) and calculate spatial correlation profiles.

In order to define characteristic spatial scales of each pattern, we used spatial correlation functions, which measure how correlated two points at a defined distance are to each other across the entire image^107^. In spatial arrangements with larger patches, high correlation is maintained for farther distances between points on average. Spatial autocorrelations of segmented (resistant/green) signal were calculated and radially averaged using the “autocorr2” function (Santiago Benito, Matlab Central File Exchange, now replaced with “twoPointProbability”) in Mathworks MATLAB 2022b. As a summary result to describe the “size” of spatial patterns of resistant/green signal, we also defined a correlation length as the distance at which radial correlation curves fell below 0.5.

### Live imaging of colony growth

In order to capture the dynamics of pattern formation, we also imaged time series of colonies overnight. To this end, after seeding bacteria onto the agar plates as above and letting the droplet dry for about 15-20 minutes, plates were placed into an incubation chamber surrounding the LSM 700 confocal microscope, which kept the temperature around 37°C to equilibrate for 30-60 minutes. Then, an imaging window with a cover glass coated with anti-fog solution (Don and Jon’s Antifog Extreme) to avoid condensation was placed onto the agar plate. The surrounding surface area of the plate was covered with a custom printed cover to slow down drying of the agar, and some moistened paper towels were kept inside the incubator to keep the humidity at a higher level. Using the autofocus function of the microscope software to adjust for movement due to drying of the agar, a section of the homeland at the agar surface was imaged every 10 minutes for up to 60 timepoints, although the usable timepoints frequently were fewer, due to the colony surface eventually touching the cover glass or water accumulating in the interface.

Due to movement of the agar over time, movies were stabilized by tracking the center of a colony manually over all time points and then selecting a 500x500 µm^2^ area around that center point at each time point.

While the green fluorophore mDasher was usually visible at early time points, mRudolph was not bright enough to be captured until much later in the growth process, and was thus not included in our analysis. Because the intensity of mDasher typically increased greatly over time, its brightness was adjusted individually for each time point using the Brightness/Contrast > Auto function in FIJI/ImageJ. To illustrate growth over time, representative lines through the image were “resliced” at a 1µm width in FIJI/ImageJ to create kymographs across all timepoints.

### Well-mixed vs spatial comparisons

From exponentially growing cultures of ampicillin-resistant (mDasher-labeled) and sensitive (mRudolph-labeled) og1rf strains, new mixed cultures on agar and in liquid media were set up in parallel. A mixture of specific bacterial density and resistant fraction in BHI was prepared as above and vortexed, and then a 2µl droplet of that culture added either onto a 35mm diameter agar plate or into a vial containing BHI in a 5ml volume, each containing 120µg/ml spectinomycin and different ampicillin concentrations as needed. After the droplets on the agar plates dried, agar plates were kept in an incubator at 37°C over night for about 18 hours.

Simultaneously, the liquid vials were kept at 37°C, with magnetic stir bars ensuring well-mixed conditions. The next day, all samples were removed from their optimal growth temperature at the same time and kept in the fridge until all samples were measured. For vials, the optical density of the media was measured after appropriate dilution to the linear range, tracking the dilution volumes. For agar plates, 1ml of BHI or PBS (Phosphate Buffer Saline) was added on top of the plate, the colony was carefully scraped off the agar surface with an inoculation loop and then the (as many as possible) bacteria were transferred to a tube in the liquid medium. After vortexing to break up any clumps of bacteria, OD measurements were made for those washed-off colonies as well. All OD measurements were multiplied by dilution volumes to compare overall population sizes across conditions.

In order to estimate the final resistant fraction of the population after overnight growth, 2 µl of each sample were placed in between two 1mm thick glass slides. Using a Zeiss LSM 700 confocal microscope, mDasher (488 nm excitation) and mRudolph (555 nm excitation) fluorescence, as well as transmitted light, were acquired. After stitching the tiles and z projecting the slices of each image by maximum intensity (as above), within an area of 312.58 µm x 312.58 µm (1000x1000 pixels) all cells were marked manually based on the transmitted light channel and counted in FIJI/ImageJ. Cells expressing green fluorescent protein (which had very little background signal) were counted as resistant, and their fraction of the total cell number was calculated.

### Antibiotic dose response on agar

To estimate the response of sensitive og1rf bacteria to different ampicillin concentrations on agar under our experimental conditions, which can differ from the dose response in liquid culture, we prepared fresh agar plates containing BHI, 120 µg/ml spectinomycin, and between 0 and 2 µg/ml ampicillin. For exponentially growing cultures of og1rf marked with mRudolph (as used in our other experiments), the density was determined, and dilutions to goal ODs of 0.1 and 0.005 (to represent the range of densities used in the main experiment) were prepared.

One 2 µl droplet was added to each agar plate, dried, and let grow at 37°C over night for about 18 hours. Then, as above, 1 ml of PBS was added to each plate, bacteria loosened from the surface with an inoculation loop and transferred in the PBS to a fresh tube. Those samples were vortexed, diluted if necessary, and the final OD determined. From 3 replicate measurements, and the mean normalized ampicillin dose response curve was fit to a hill-like function 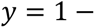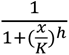 (Figure S9A). From the OD 0.005 condition, parameters K = 0.24 and h = 3.22 were estimated and used to inform model parameters. From the OD 0.1 condition, parameters of K = 0.20 and h = 1.74 were fitted. The difference in the Hill coefficient h fit (and poor Hill fit at OD 0.1) is potentially caused by a “bump” in the OD 0.1 dose-response curve around 0.5- 0.75 µg/ml Ampicillin, deviating from a sigmoid shape (Fig. S9A), possibly due to community effects on resistance that would be interesting to investigate in the future. The same experiment was conducted with freshly dissolved amoxicillin instead of ampicillin to measure dose response curves for amoxicillin (Figure 5C).

### Sulbactam experiments

To test how well our model predicted the effect of slowing down β-lactamase activity, we repeated our agar colony experiment while combining different concentrations of ampicillin and the β-lactamase inhibitor sulbactam (Alfa Aesar/Thermo Fisher Scientific, 233.24 g/mol ), keeping all other conditions the same.

### Computational model

Simulation scripts were run in Matlab 2022b and can be accessed at 10.5281/zenodo.14226311. To model growth of resistant and sensitive bacteria on a surface, we implemented a 2D cellular automaton model coupled to a reaction-diffusion model of antibiotic degradation. The bacterial growth was simulated on a 500x500 grid, representing a ∼500x500 µm^2^ patch of the homeland with periodic boundaries. Periodic boundaries were chosen in order to simulate a subregion of the seeded area, neighbored by similar subregions. However, the model results were similar when no-flux boundaries were applied (Figure S21). In this model, each grid position corresponded to space for a single bacterium (∼1 µm diameter), and “resistant“ and “sensitive” initial grid positions were assigned randomly at fixed ratios corresponding to experimental conditions. The total number of initial cells was set to 182 for OD 0.01, 467 for OD 0.03 and 1443 for OD 0.1, based on an experiment counting bacteria in a 500x500 µm^2^ area of the homeland right after seeding (2 replicates, Figure S22). (These densities are lower than expected from total estimated cell number based on OD and droplet area, possibly due in part to the coffee ring effect ^108,109^.

Ampicillin diffusion speed was estimated roughly based on measurements for penicillin diffusion through agar ^110^ (0.016 cm^2^/hr = 2.67 x 10^5^ µm^2^/growth timestep). Diffusion was modeled using a forward time central space (FTCS) scheme of convolution with directly neighboring pixels on a 3D 500 µm x500 µm x500 µm (100x100x100 voxels) grid, the top layer of which was where the 2D cell grid was located, using the MATLAB imfilter function and based on the simulation used in ^74^ . The boundaries of the antibiotic grid were periodic in x and y directions, matching the cell grid, but reflective in z. In order to make the simulation more feasible, antibiotic diffusion was coarse grained to 5 µm x 5 µm x 5 µm voxel sizes, creating the 100x100x100 grid (diffusion per timestep was limited due to the one-pixel distance of convolution for stability: → 0.15 → for 1µm voxels ∼1’800’000 iterations per growth timestep → coarse grained voxels → iterations divided by 5x5 - 720’000).

Antibiotic concentration was set uniform across the whole grid to a global concentration between 0 and 2 (equivalent of µg/ml based on sensitive death rate) at the beginning of the simulation and after each growth time step. Then, changes to the antibiotic landscape were calculated from local degradation at locations occupied by resistant bacteria and diffusion of antibiotic throughout the 3D grid. After the 720’000 diffusion and degradation iterations corresponding to ∼10 minutes, the resulting antibiotic distribution was used to calculate local death probabilities for the next growth step. Local degradation by resistant cells in the top layer was assumed to be very fast ^98^ even relative to diffusion, such that the antibiotic concentration at each position occupied by resistant cells was set to 0 at each diffusion time step. Consistent with previous experimental results, we assumed that β-lactamase stayed bound to the resistant cells in *E. faecalis* ^62,63^ and thus only considered degradation at the locations of resistant cells.

After calculating the antibiotic distribution based on degradation and diffusion, growth and death decisions were calculated based on randomized decisions for each cell, and the cell grid was updated.

Growth was assumed to have equal baseline rates for both cell types and be independent of antibiotic. The probability of a birth is decided for each individual position r based on the following:

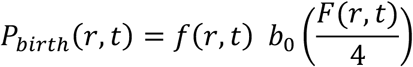

Where f(r, t) is the occupancy of that position (0 if empty, 1 if full) at time t, b_0_ is the base growth rate (the same for resistant and sensitive bacteria), and F (t, r) is the current number of free nearest (Von Neumann range 1) neighbors (between 0 and 4) – that determines the competition effect. Growth rates b_0_ were set so that on average half of bacteria divided at each time step. Thus, assuming a division time of about 20 minutes ^111^, each time step roughly represents 10 minutes.

The probability of death for each individual position is determined by:

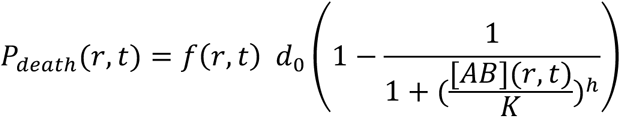

Where f(r, t) is the occupancy of that position (0 if empty, 1 if full), d_0_ is the base death rate (0 for resistant bacteria and equal to b_0_ for sensitive bacteria – this relates the two equations so if [AB] equals the minimum inhibitory concentration, sensitive death and birth rates would be equal in absence of competition), [AB] (r, t) is the current antibiotic concentration at the equivalent location in the grid, and K and h are Hill function coefficients (K = 0.325 and h = 1) obtained by approximating experimental dose response curves of sensitive bacteria (measured as described above). We empirically fit the dose response parameters by running several sensitive-only simulations and aiming for similar final sensitive densities as in the experiment (Figure S9A). The model did not include resistant cell death, because in our experimental results significant resistant death appeared to only occur at very low resistant cell densities and the highest antibiotic concentrations used (Figures 1 and S1-3).

In this model, only cells at open surfaces are able to divide. This is a common assumption in, e.g. classical Eden growth models ^73^ or range expansion models, since bacteria behind the front often stop growing (at least in the two-dimensional plane) either due to low nutrient access or physical pressure ^82,84^. We initially tried to match the total time of simulation to the experiment (∼108 10-minute timesteps for 10 hours), but this did not allow the simulations to fill in the area to a comparable extent as seen in the experiment (Figure S23A). Comparing our experimental and simulation timeseries suggests that while the patch radius expands approximately linearly in the simulation before encountering competition, experimental clusters expand at a changing rate that accelerates (Figure S23B). Using a less restrictive model that allows new cells to “jump” up to three grid steps from their mother cell showed that the slow growth in the simulation can at least in part be explained by the competitive restriction of growth (Figure S23C).To adjust for this difference, we scaled the simulation time to 400 timesteps, after which the amount of area coverage was similar to the experiment (Figure S23A). The antibiotic surface and resistant and sensitive grids were saved every 5^th^ timestep, to track the evolution of the simulation over time. For both experiments and simulations, patterns and composition reached a state of relative stability after the surface area was filled (Figures S7 and S14, respectively).

The depth of the antibiotic 3D simulation was set to 500 µm to balance simulating a large global supply of antibiotic relative to the colony size, and keeping the simulation time and memory requirements reasonable. We also tested on a smaller xy grid that increasing the depth of the simulation did not have a large effect on the resulting patterns (Figure S24).

For OD 0.01 and OD 0.03, a resistant:sensitive ratio of 0.1:99.9 resulted in no resistant cells being seeded at the grid size we had chosen for feasibility reasons. Those simulations were thus done separately on an 1250x1250 grid.

Simulation results were largely consistent across replicates for representative cases (Figure S25) in terms of both composition and pattern scale, despite the stochastic nature of the simulation. Thus, we deemed single runs sufficient for all other homeland simulations, especially considering the long run time of the simulation.

### Non-cooperative simulation

To test to what extent cooperation by active antibiotic degradation plays a role in pattern formation, we also ran a simulation in which antibiotic was not degraded and was thus uniformly maintained at the global initial concentration across the entire grid (Figure S9B).

### Sulbactam and amoxicillin simulations

To simulate β-lactamase being slowed down by an inhibitor, we ran simulations in which degradation only occurred every 5^th^ or 10^th^ timestep, rather than every timestep as in the original simulation. We did not have a clear way to measure by how much sulbactam slows down β-lactamase activity in the experiment, so we just identified what range of degradation frequencies matched the experimental data, and maintained analogous linear scaling (i.e. if 2 µg/ml sulbactam correspond to degradation every 10^th^ step, 1 µg/ml corresponds to degradation every 5^th^ step).

To predict the effect of amoxicillin on homeland patterns, we used the dose response curve of sensitive cells (see above) measured in a separate experiment and adjusted the death rate parameters in our model accordingly. We then conducted experiments combining resistant and sensitive cells to compare to our model predictions.

### Range expansion simulations

To investigate the effect of degradation in the homeland on subsequent range expansion, we modified our simulation to randomly seed a circle with a radius of 74 µm and a width of 2µm at specified resistant:sensitive ratio, to simulate the growth front of range expansion. The inside of the circle was then filled either with a random distribution of resistant and sensitive cells at the same ratio (simulating a densely seeded colony), or with non-growing cells that either were all non-degrading, or all degrading antibiotic (Figure 5B). Otherwise the model stayed the same as the main simulations.

The simulation was then let evolve for 200 timepoints, which was sufficient to observe a broad band of range expansion (∼40 µm width). We did 3 replicates for this simulation since range expansion appeared more sensitive to stochastic events than homeland patterning (Figure S16). We then analyzed the composition across the range expanding region using a radial average at equal distances from the center of the homeland. To reduce noise, a 5 µm moving average smoothing was applied to all curves before combining the three replicates.

## Acknowledgments

We thank the Wood lab, as well as A. A. King, L. Zaman, R. J. Woods, A. C. Martin and S. Veatch, for helpful discussions throughout the writing and revision process, and the two anonymous reviewers for thoughtful comments that have improved our manuscript. This work was supported by the National Institutes of Health (NIH) Award R35GM124875 to the University of Michigan, awarded to K. B. W., and a fellowship from the Jane Coffin Childs Memorial Fund for Medical Research, awarded to M. K. D.-L.

See Supplemental Information for Supplemental Figures and Movie Legends.

## Notes

https://doi.org/10.5281/zenodo.14226310

### Competing Interest Statement

The authors have declared no competing interest.

### Summary of Updates

Refocused the framing to more accurately reflect the contributions of this work, added explanation and controls for the biophysical model, improved some figure readability concerns.

