## Supplementary Figures and Movie Legends for "Spatial population dynamics of bacterial colonies with social antibiotic resistance"

**Supplemental Information for Denk-Lobnig and Wood, "Spatial population dynamics of bacterial colonies with social antibiotic resistance"**

Supplemental Figures:

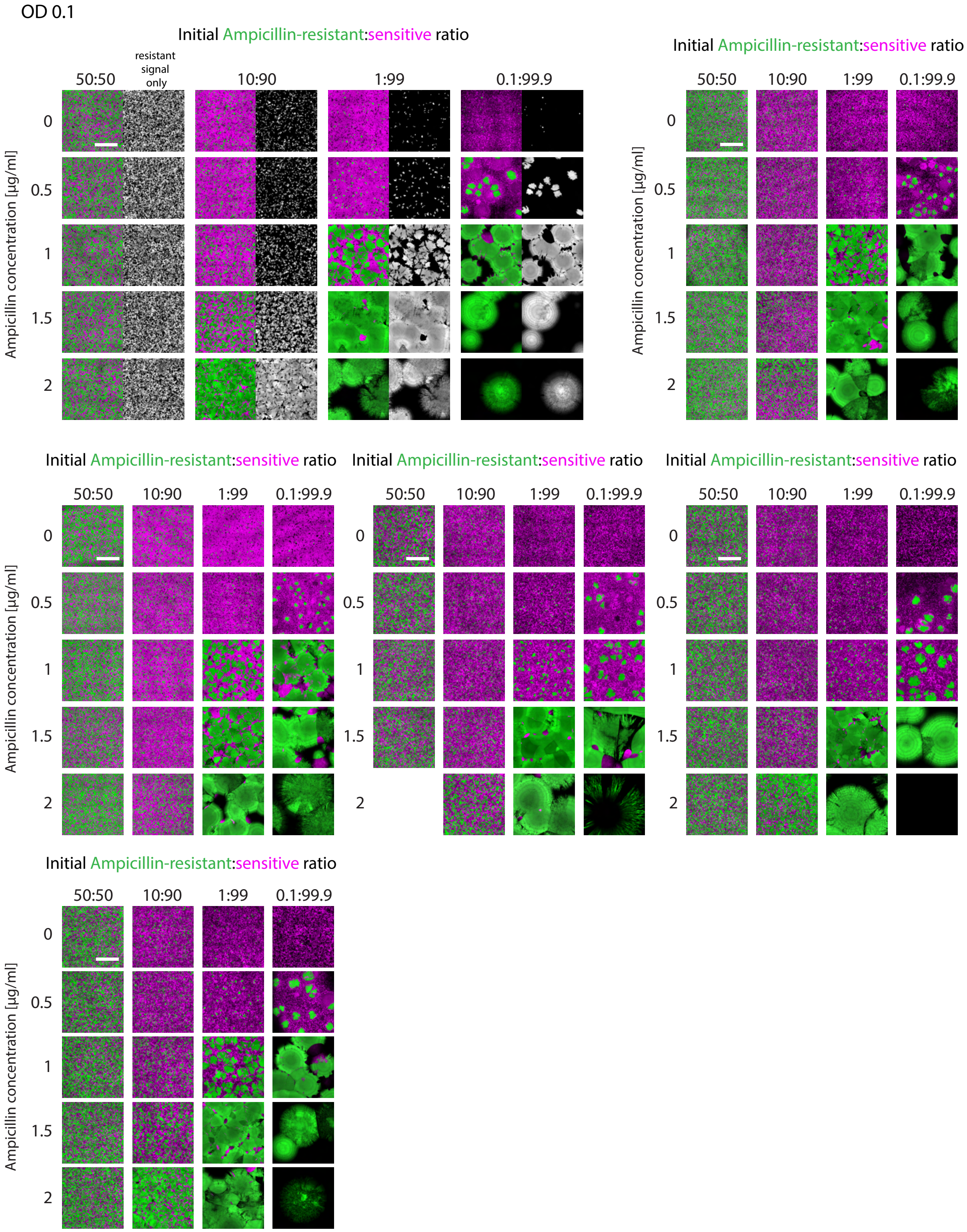


**Figure S1: Experimental replicates with an initial density of OD 0.1.** Confocal images showing the homeland of colonies across all replicates that started with an OD of 0.1, for the main dataset shown In Figure 1 and analyzed in Figure 2. Each grouped panel represents colonies with different initial conditions that were imaged within the same experiment or same week. The top left panel shows images combining resistant (green) and sensitive (magenta) channels, as well as a corresponding gray scale images of the “resistant” channel (488 nm), for the replicate also shown in Figure 1. Missing panels represent samples that were not suitable for imaging and analysis due to issues in the experimental process. Scale bars = 500 nm, all images shown are at the same scale.

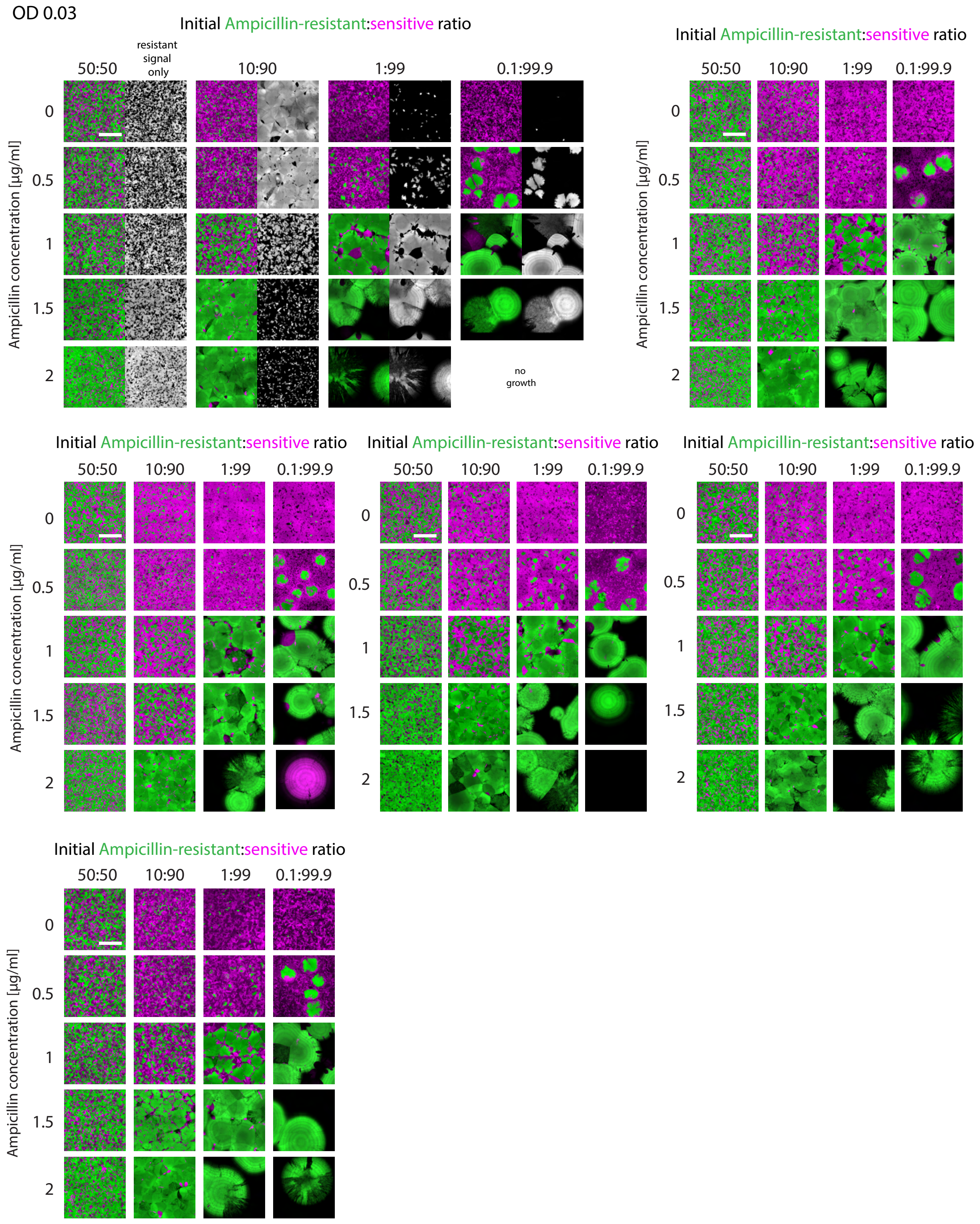


**Figure S2: Experimental replicates with an initial density of OD 0.03.** Confocal images showing the homeland of colonies across all replicates that started with an OD of 0.03, for the main dataset shown In Figure 1 and analyzed in Figure 2. Each grouped panel represents colonies with different initial conditions that were imaged within the same experiment or week. The top left panel shows images combining resistant (green) and sensitive (magenta) channels, as well as corresponding gray scale images of the “resistant” channel (488 nm), for the replicate also shown in Figure 1. Missing panels represent samples that were not suitable for imaging due to issues in the experimental process or where no growth occurred. Scale bars = 500 nm, all images shown are at the same scale.

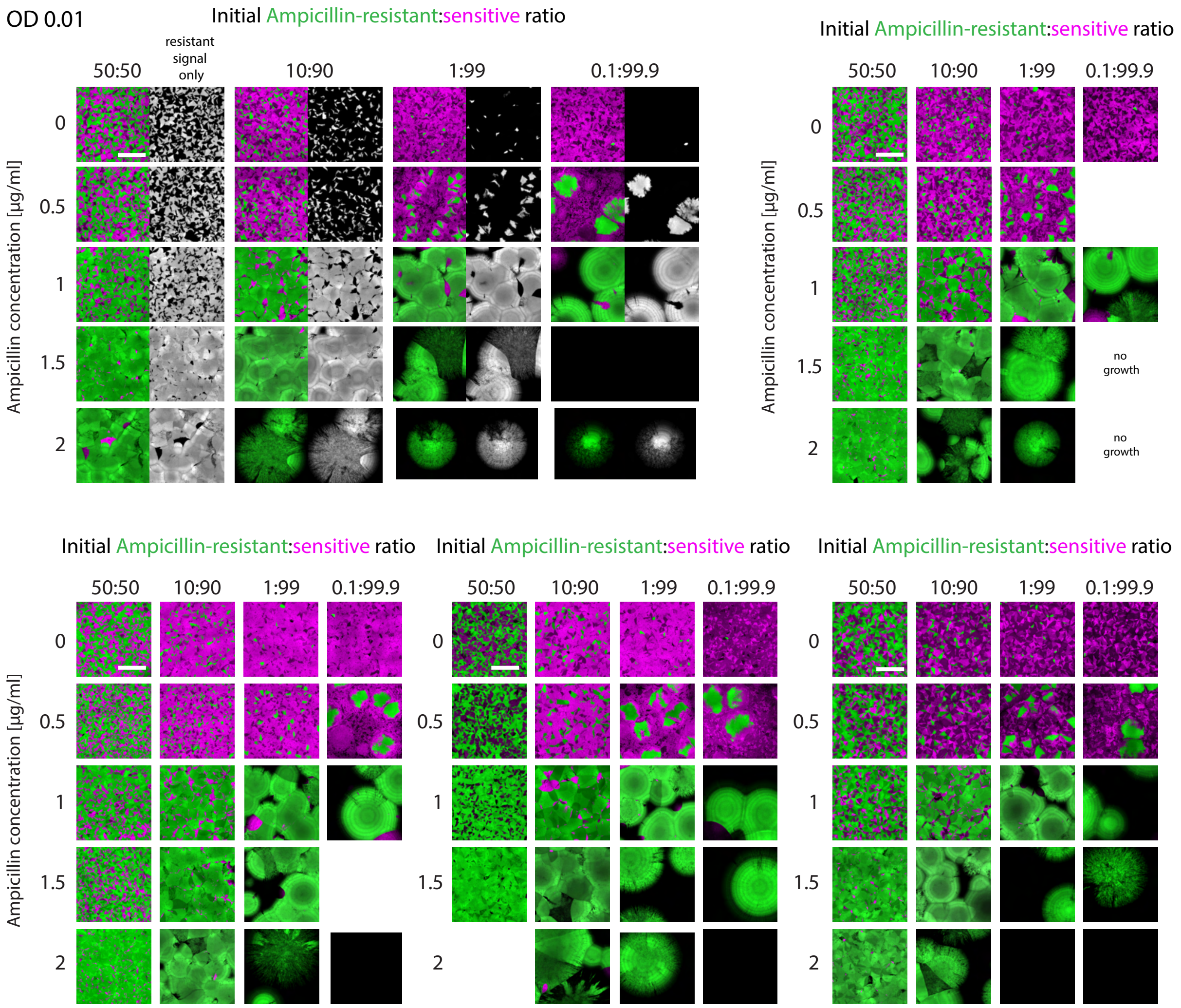


**Figure S3: Experimental replicates with an initial density of OD 0.01:**
Confocal images showing the homeland of colonies across all replicates that started with an OD of 0.01, for the main dataset shown In Figure 1 and analyzed in Figure 2. Each grouped panel represents colonies with different initial conditions that were imaged within the same experiment or the same week. The top left panel shows images combining resistant (green) and sensitive (magenta) channels, as well as corresponding gray scale images of the "resistant" channel (488 nm), for the replicate also shown in Figure 1. Missing panels represent samples that were not suitable for imaging due to issues in the experimental process or where no growth occurred. Scale bars = 500 nm, all images shown are at the same scale.

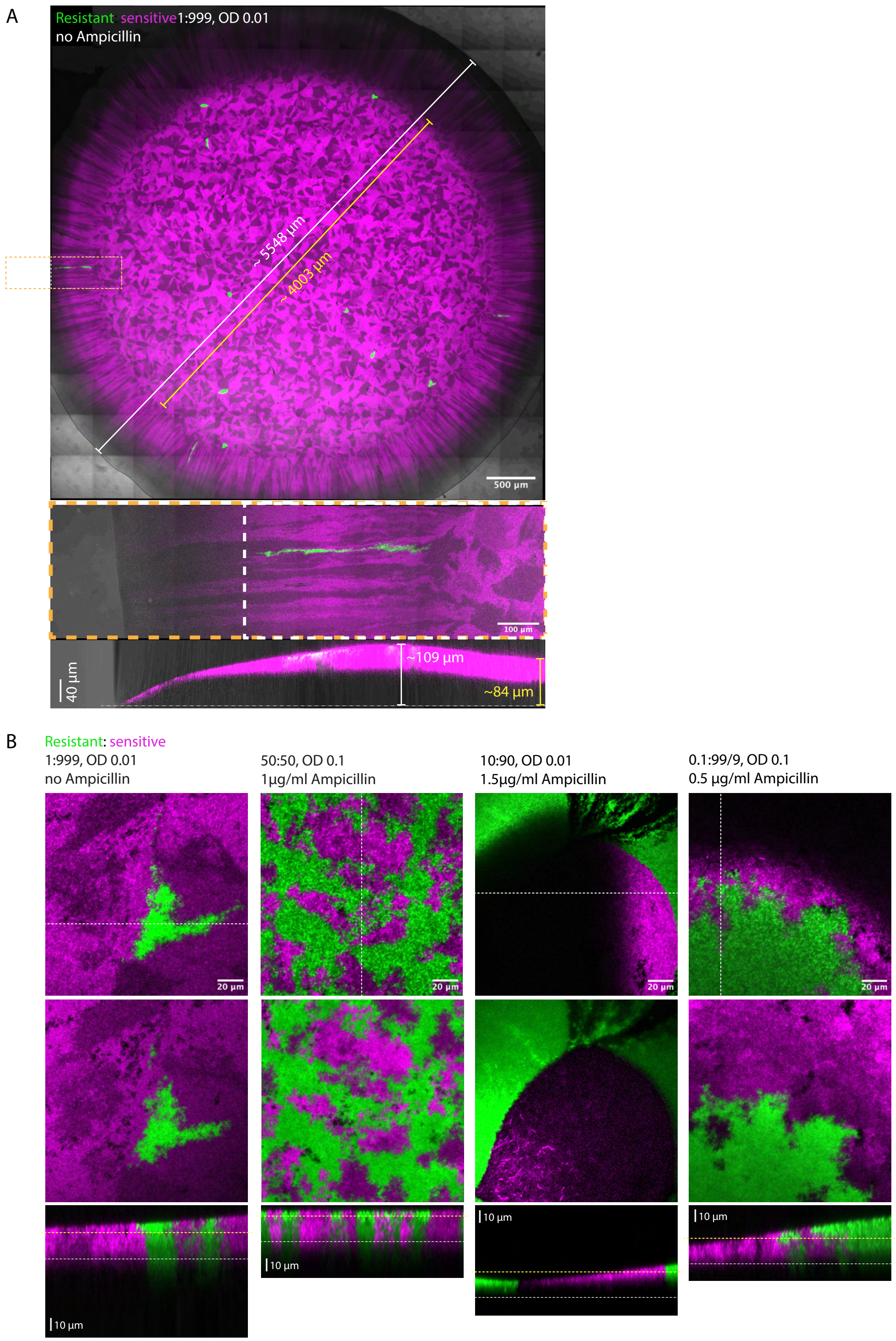


**Figure S4: Dimensions of E. faecalis colonies and pattern consistency across colony depth.** (A) Top: Maximum intensity projection of confocal microscopy image showing an *E. faecalis* colony grown for about 18 hours at 37˚C on agar. Resistant strain fluorescent signal in green, sensitive in magenta, transmitted light in grayscale. White and yellow lines and text show the diameter of the entire colony and the (approximate) homeland, respectively. Dashed yellow rectangle shows area also displayed in the images below, acquired separately at higher resolution. Top view maximum intensity projection (middle) and cross-sectional view (bottom) of the colony edge, indicating the colony height within the homeland and at the homeland-range expansion boundary in yellow and white, respectively. Scale bars as labeled. (B) Single confocal images at the surface (top) and ~20µm below the surface (middle) of *E. faecalis* colonies, as well as cross-sectional views (bottom), across several different initial conditions (OD = optical density at 600 nm wavelength). Resistant strain in green, sensitive strain in magenta. White dashed lines in top images represent the position of the cross-section; yellow and white lines in the bottom image indicate z-positions of the individual slices shown above. Scale bars as labeled.

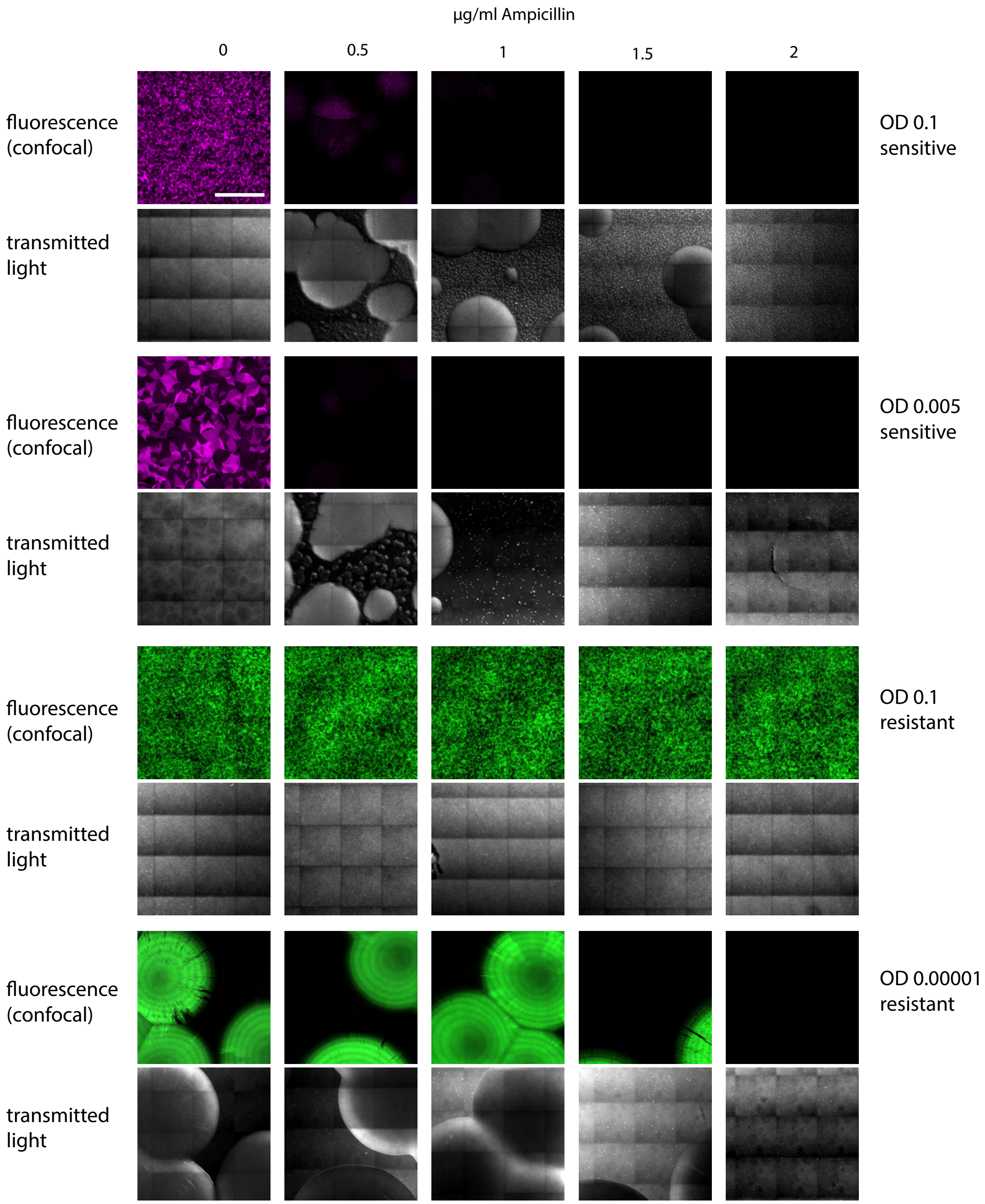


**Figure S5: Colonies with only ampicillin-sensitive or only ampicillin-resistant populations at different initial densities.** Confocal fluorescent and transmitted light maximum intensity z-projections of the more ampicillin-sensitive mRudolph strain (magenta) and the β-lactamase expressing, more ampicillin-resistant mDasher strain, across two different initial optical densities (OD), exposed to different ampicillin concentrations in the agar and grown ~18 hours at 37˚C. The shown densities correspond to the maximum and minimum individual densities of each strain across the main dataset (Figure 1). For the sensitive strain, a clear reduction in biomass in fluorescence and growth is visible with increasing ampicillin concentration at both initial densities. For the resistant strain, an inoculum effect is visible, as at the low initial density there is no growth at 2µg/ml ampicillin, whereas the high initial density condition grows seemingly without perturbation by the antibiotic. Natural variability in fluorescence intensity across the colony is visible with or without ampicillin, in addition to some uneven illumination effects at the edges of individual tiles of the tile scan. Scale bars 500 µm.

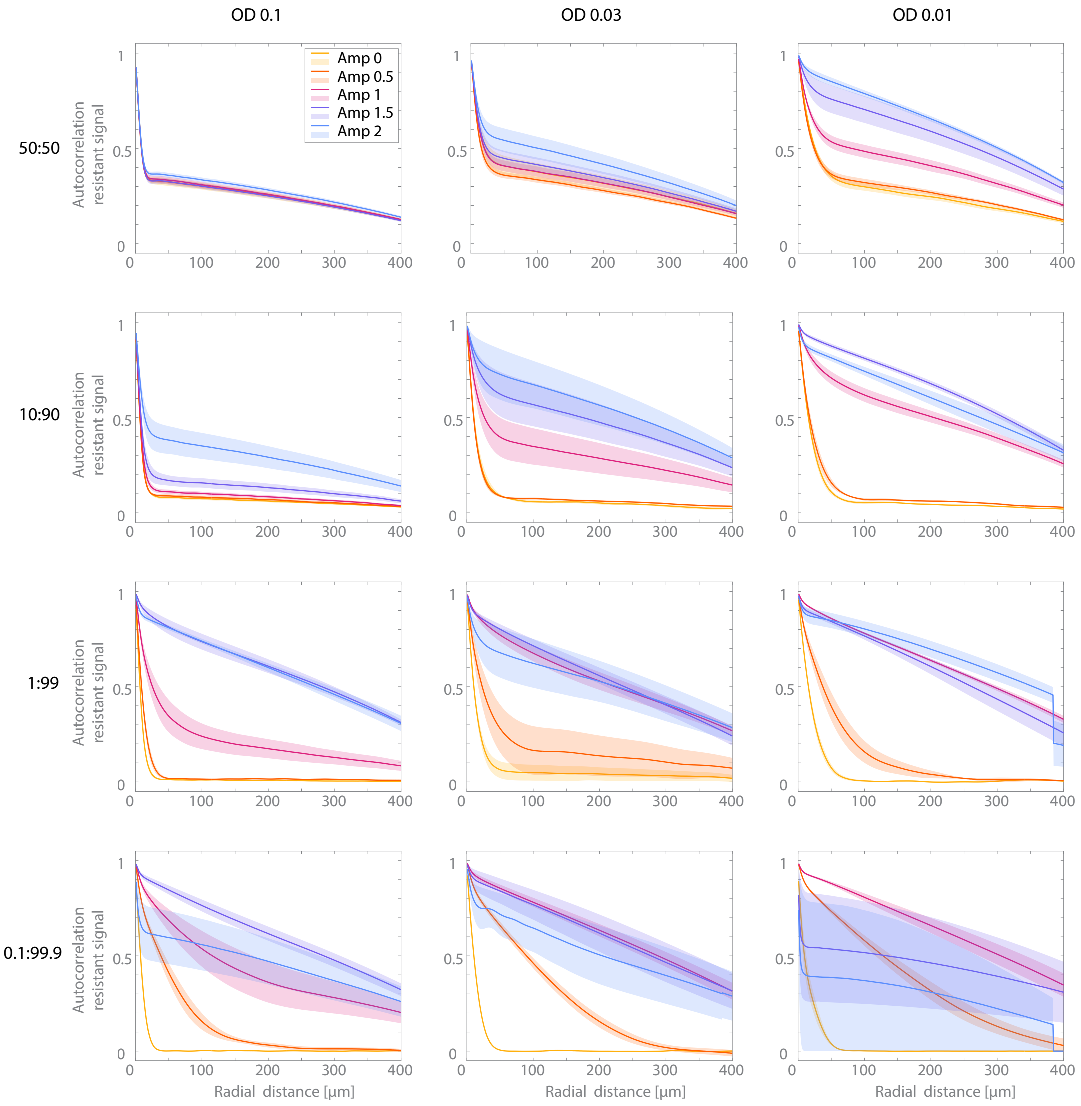


**Figure S6: Radially averaged resistant autocorrelations illustrate the relationships between initial conditions and pattern size.** Full dataset (including examples shown in Figure 2B and the basis for summary data in Figure 2C) showing radially averaged spatial correlation functions for the segmented green (resistant) fluorescent signal across all initial conditions. Mean lines and standard error as shaded area are displayed (n=3-6 for each condition, see Figures S1-3 ). Across most initial conditions, higher Ampicillin concentrations cause autocorrelation functions to decline more slowly, indicating an increase in average resistant patch size or clustering. This effect is more pronounced as the overall density or resistant density decreases. For high ampicillin concentrations paired with very low resistant fractions, autocorrelations tend to have higher variability or a faster declining average because the growth and survival of resistant cells start being affected under these conditions.

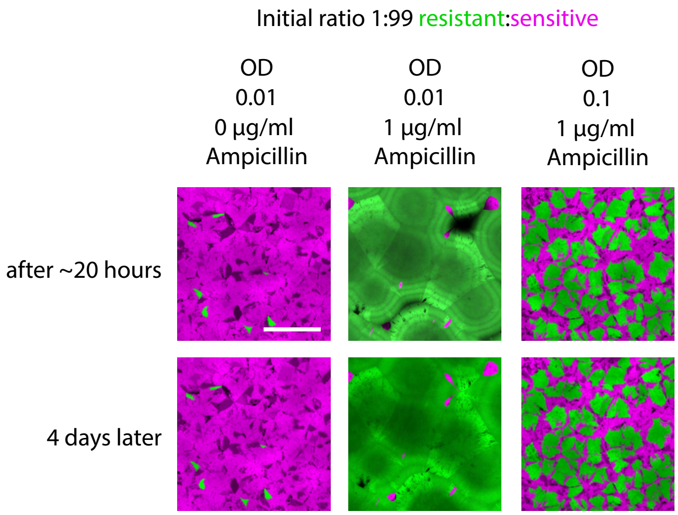


**Figure S7: Colonies imaged after overnight growth and again after 3 additional days maintain stable spatial organization.** Confocal images of colonies after ~18 hours of growth at 37˚C (top) and of the same colonies after another 4 days of growth at 37˚C, for three different initial conditions. Resistant strain in green, sensitive strain in magenta. Scale bar 500 µm.

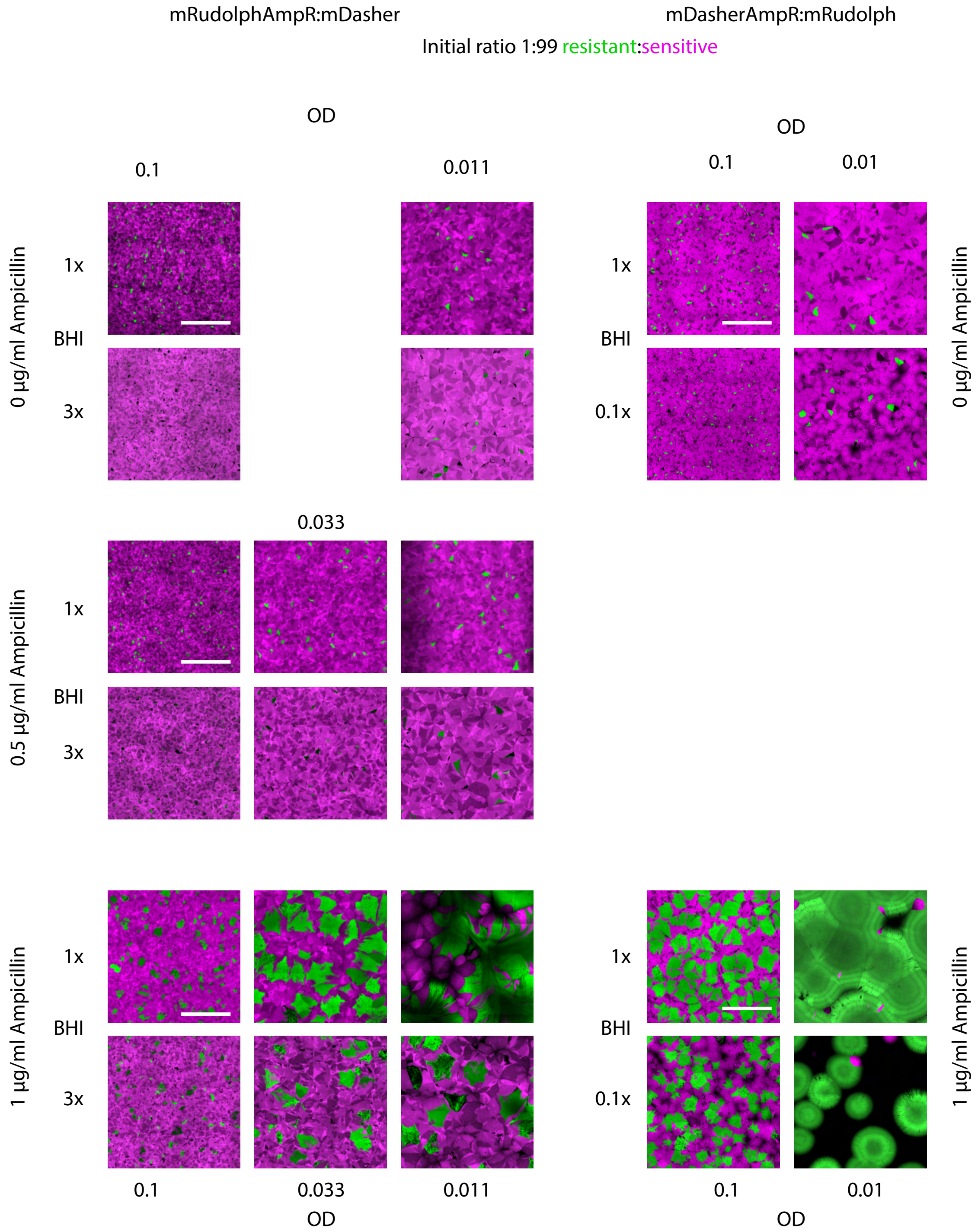


**Figure S8: Effects of changing nutrient (media) concentration on spatial organization under antibiotic exposure.** Confocal images showing colonies grown on agar containing either the regular concentration of Brain Heart Infusion media (BHI) (top rows, respectively), or either 3-fold (left) or 1/10$^{th}$ (right) of that concentration (bottom rows, respectively), across different initial densities and ampicillin concentrations. Images from high BHI conditions sometimes appear brighter because of autofluorescence of the media. Resistant bacteria are shown in green (on the left, the mRudolph AmpR strain, imaged at 555 nm, on the right, the mDasher-AmpR strain, imaged at 488 nm) and sensitive bacteria are shown in magenta (on the left, the mDasher strain, imaged at 488 nm, on the right, the mRudolph strain, imaged at 444 nm). Scale bars = 500 µm.

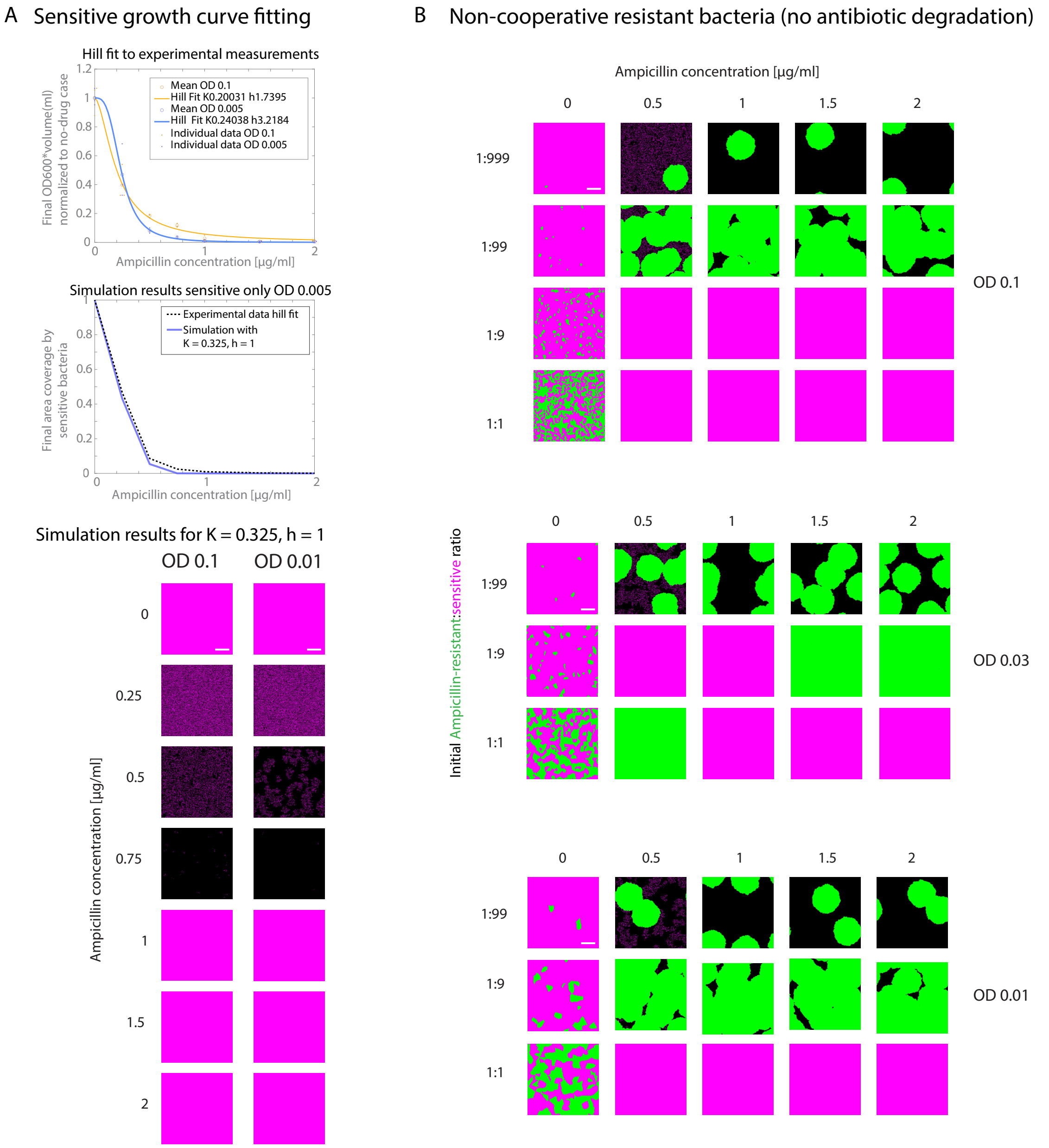

**Figure S9: Death rate parameter fitting and the importance of cooperation in the model.** (A) Top: Experimental density measurements of sensitive-only colonies washed off agar plates with different ampicillin concentrations after 18h of growth at 37˚C, at two initial densities (n = 3 for each), as well as best fit hill-like functions for each density. Middle: Hill curve from experimental fit to OD 0.005, as well as simulation final densities after 400 timesteps for a sensitive only simulation with initial OD 0.005 equivalent (91 cells), with death curve parameters used for all simulations. Bottom: Simulation of sensitive only colonies (positions occupied by cells in magenta) at two different initial densities for the same death curve parameters, across different ampicillin concentrations. Scale bars = 100 µm. (B) Simulation results without degradation of antibiotic (but only sensitive cells experience antibiotic-induced death), which simulates non-cooperative resistance and does not match experimental data for equivalent concentrations well (see Figure 1). Resistant green, sensitive magenta, scale bars = 100 µm.

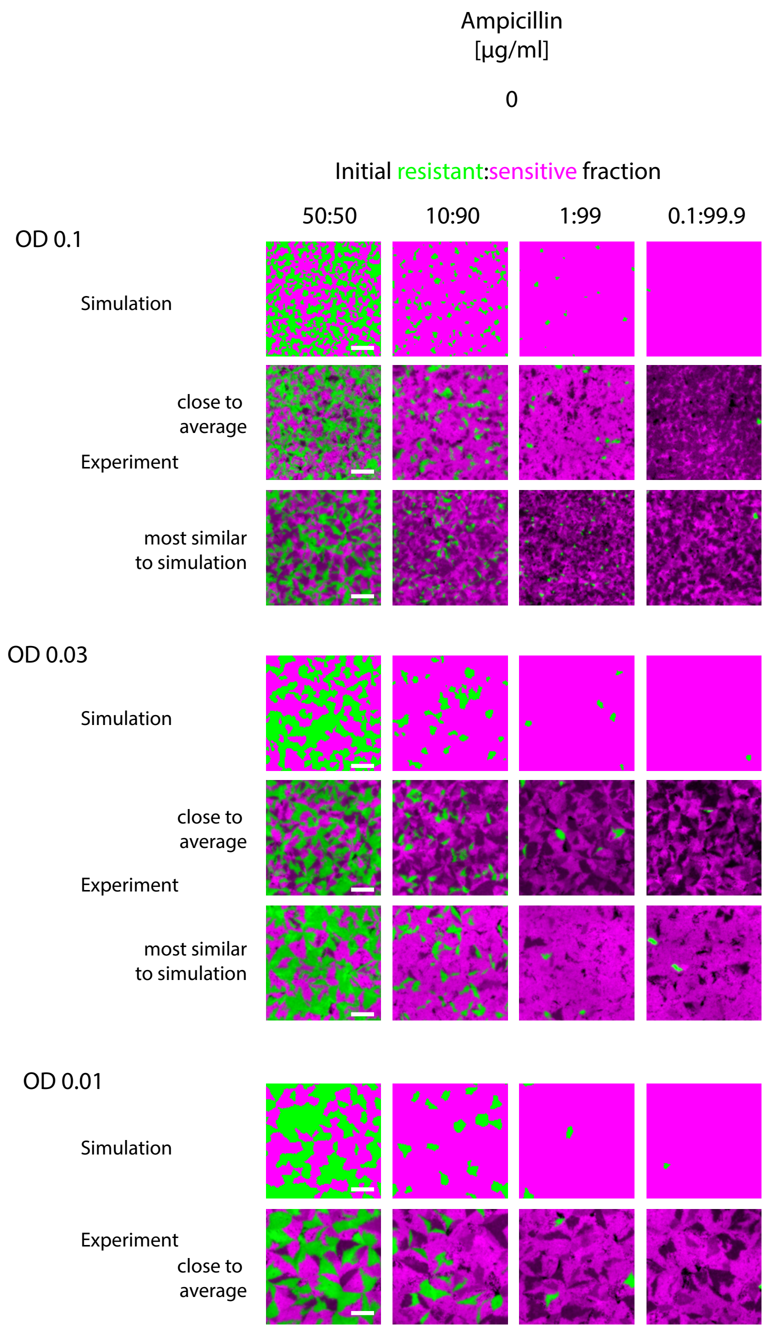


**Figure S10: Comparison of simulated and experimental patterns at 0 µg/ml Ampicillin.** Simulation results without ampicillin (top rows), 500 µmx500 µm regions of corresponding most representative replicate dataset of experimental data (middle rows, same as Figure 1) and (for OD 0.1 and 0.03) an experimental replicate appearing more similar to simulation results on average (bottom rows, to show that the simulation at times falls close to the range of experimental outcomes, even when it is not as similar to the average result), across all initial densities and resistant:sensitive ratios. Resistant in green, sensitive in magenta. Scale bars = 100 µm. For OD 0.03 and OD 0.01 at 0.1:99.9 resistant:sensitive, a representative 500 µm x 500 µm area of a 1250 µm x 1250 µm simulation grid is shown. Some examples also shown in Figure 3.

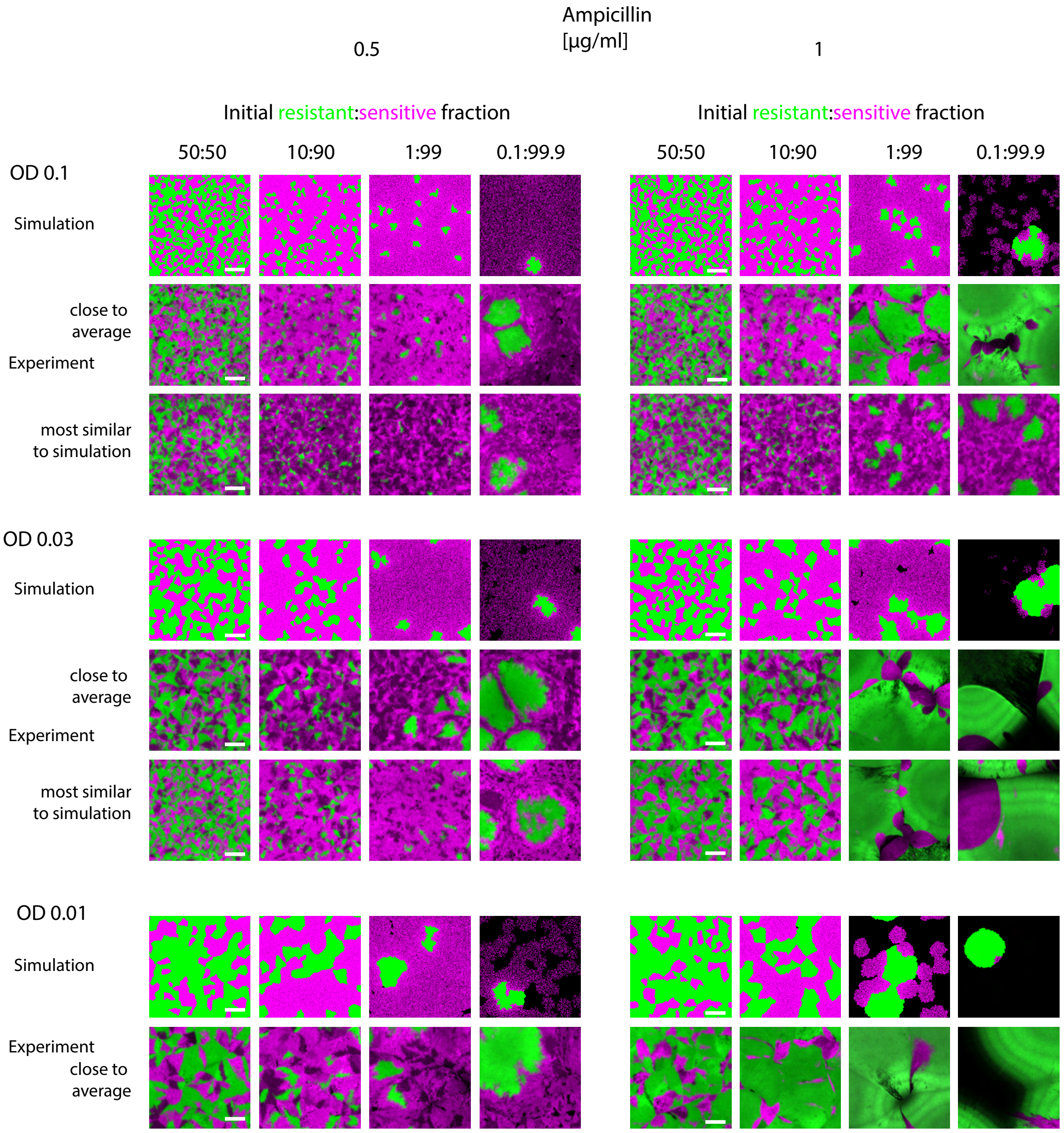


**Figure S11: Comparison of simulated and experimental patterns at 0.5 and 1 µg/ml Ampicillin.** Simulation results with 0.5 (left) and 1 (right) µg/ml ampicillin (top rows), 500 µmx500 µm regions of corresponding most representative replicate dataset of experimental data (middle, same as Figure 1) and (for OD 0.1 and 0.03) an experimental replicate appearing more similar to simulation results on average (bottom, to show that the simulation at times falls close to the range of experimental outcomes even when it is off the average result), across all initial densities and resistant:sensitive ratios. Resistant in green, sensitive in magenta. Scale bars = 100 µm. For OD 0.03 and OD 0.01 at 0.1:99.9 resistant:sensitive, a representative 500 µm x 500 µm area of a 1250 µm x 1250 µm simulation grid is shown.  Some examples also shown in Figure 3.

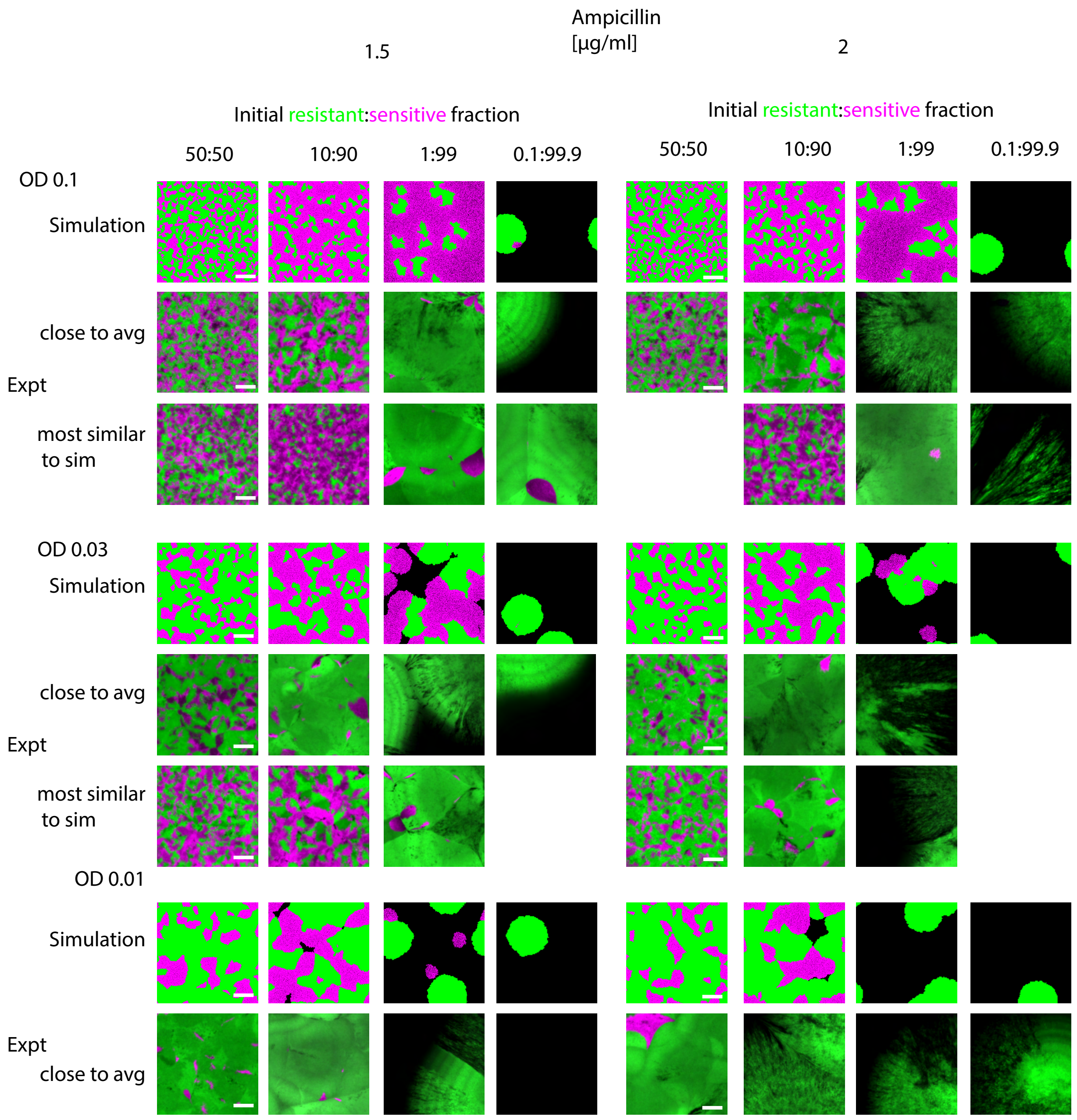


**Figure S12: Comparison of simulated and experimental patterns at 1.5 and 2 µg/ml Ampicillin.** Simulation results with 1.5 (left) and 2 (right) µg/ml ampicillin (top rows), 500 µmx500 µm regions of corresponding most representative replicate dataset of experimental data (middle, same as Figure 1) and (for OD 0.1 and 0.03) an experimental replicate appearing more similar to simulation results on average (bottom, to show that the simulation at times falls close to the range of experimental outcomes even when it is off the average result), across all initial densities and resistant:sensitive ratios. Resistant in green, sensitive in magenta. Scale bars = 100 µm. For OD 0.03 and OD 0.01 at 0.1:99.9 resistant:sensitive, a representative 500 µm x 500 µm area of a 1250 µm x 1250 µm simulation grid is shown. Some examples also shown in Figure 3.

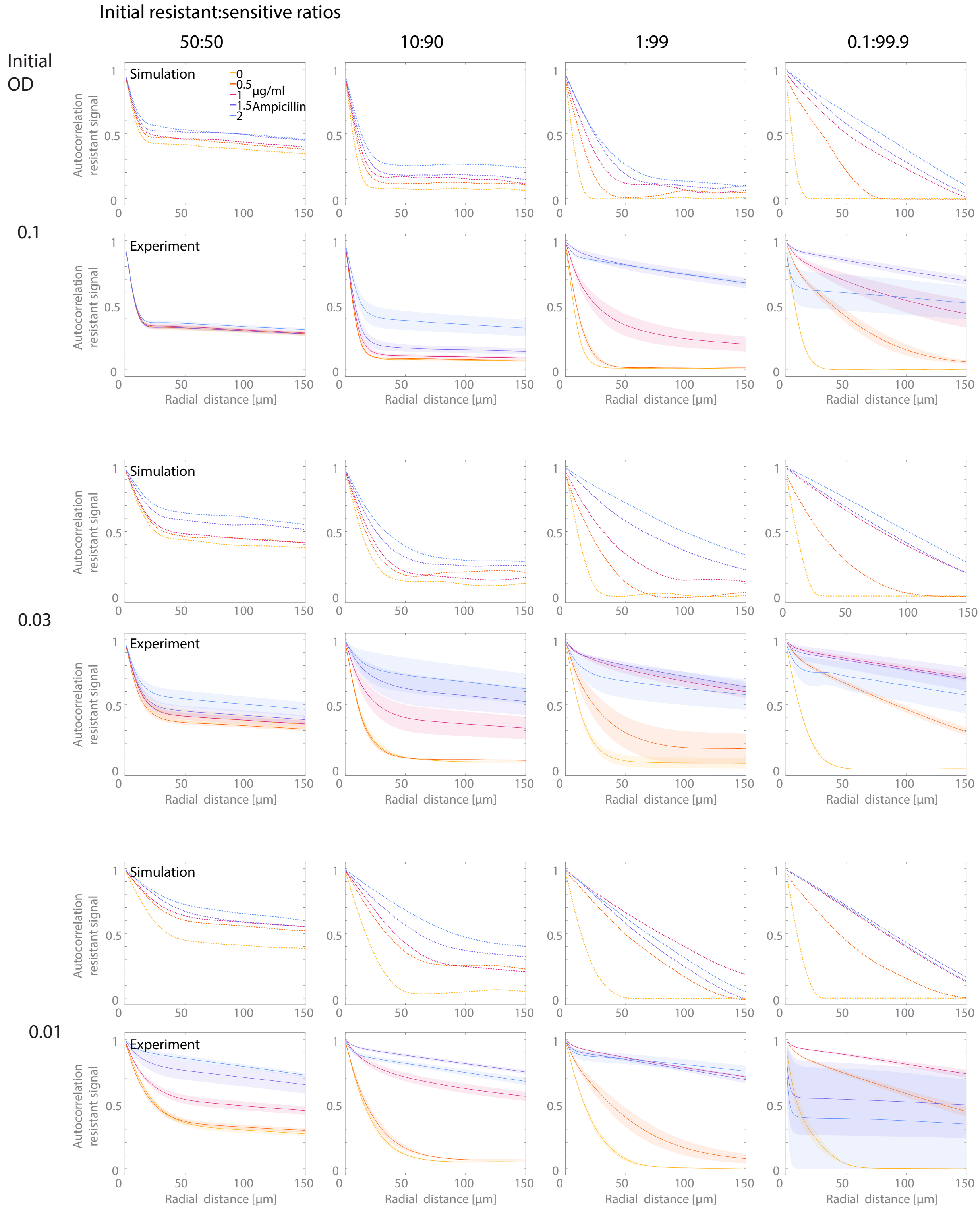


**Figure S13: Comparison of radially averaged autocorrelation functions of green (resistant) signal for simulations vs experimental data.** Full dataset (including examples shown in Figure

3E, experimental data same as Figure S6) showing radially averaged spatial correlation functions for the resistant signal in images obtained by simulations and experiments (mean lines and standard error as shaded), respectively, as a measure of pattern size for all tested initial conditions. (n = 1 for simulations, n = 3-6 for experiments). The correlation distance displayed on the x-axis is shorter than in Figures 2 and S6, because simulation grids were smaller (500 µm x 500 µm) than the analyzed homeland regions in the experiments (1339.65 µm x 1339.65 µm). Overall, while there is not always an exact quantitative agreement between model and experimental autocorrelation functions for the same condition, the trends for how autocorrelation and spatial scale vary with initial conditions are largely consistent between model and experimental results.

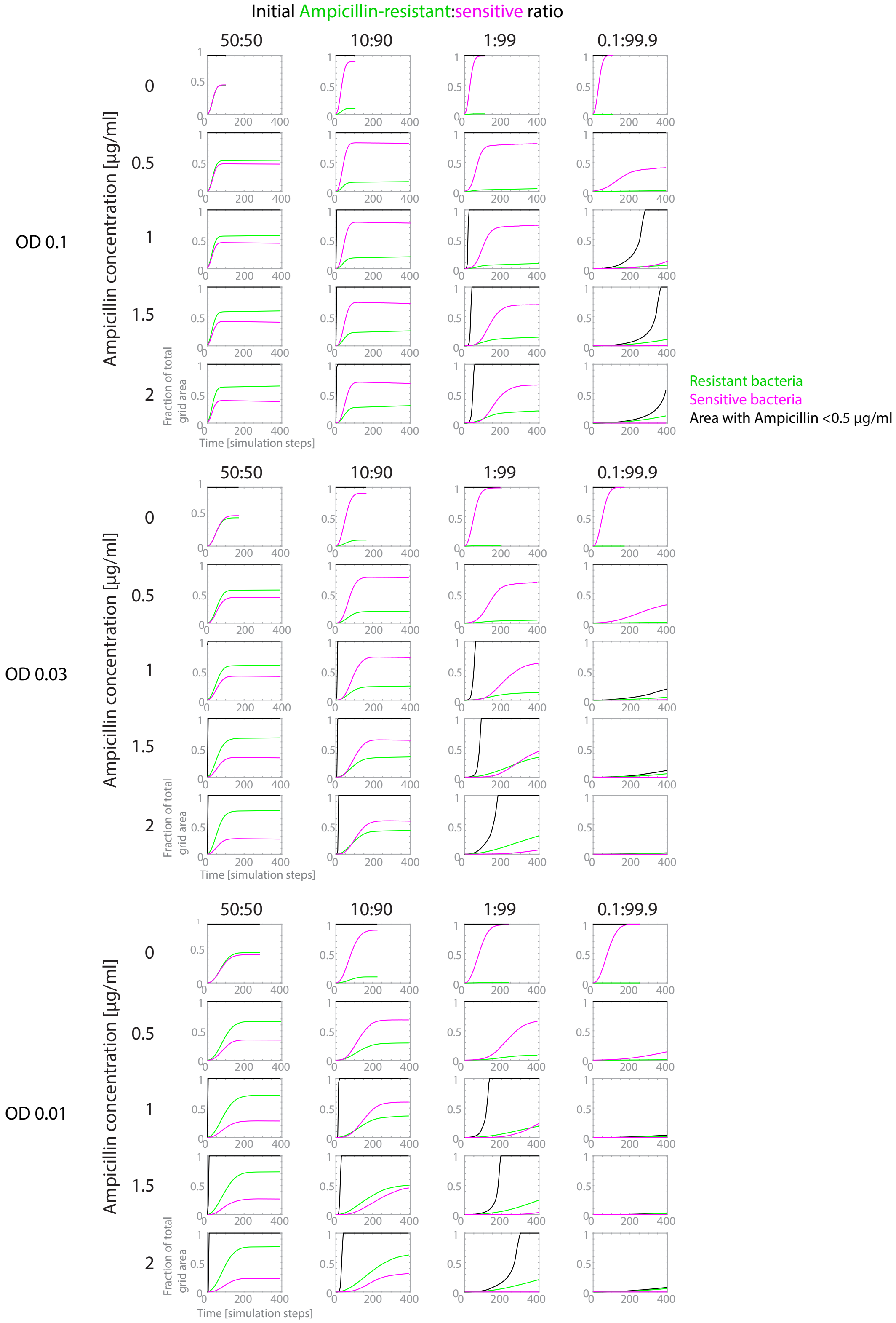


**Figure S14: Evolution of resistant and sensitive growth and antibiotic-depleted area over time in simulations of all tested conditions.** Full dataset showing simulated resistant- (green) and sensitive- (magenta) occupied and antibiotic-depleted (< 0.5 µg/ml, black) areas as a fraction of total grid area over simulation time (some examples are also shown in Figure 4) for all simulated initial conditions. The trends in this data show that the timing of a global transition from “high” (>0.5 µg/ml) to “low” (<0.5 µg/ml) drug concentration (and resulting growth or death dynamics) depends on initial drug concentration, but also initial overall population and resistant sub-population density. This suggests a mechanism for how the dynamics of growth result in differences in final composition and spatial organization.

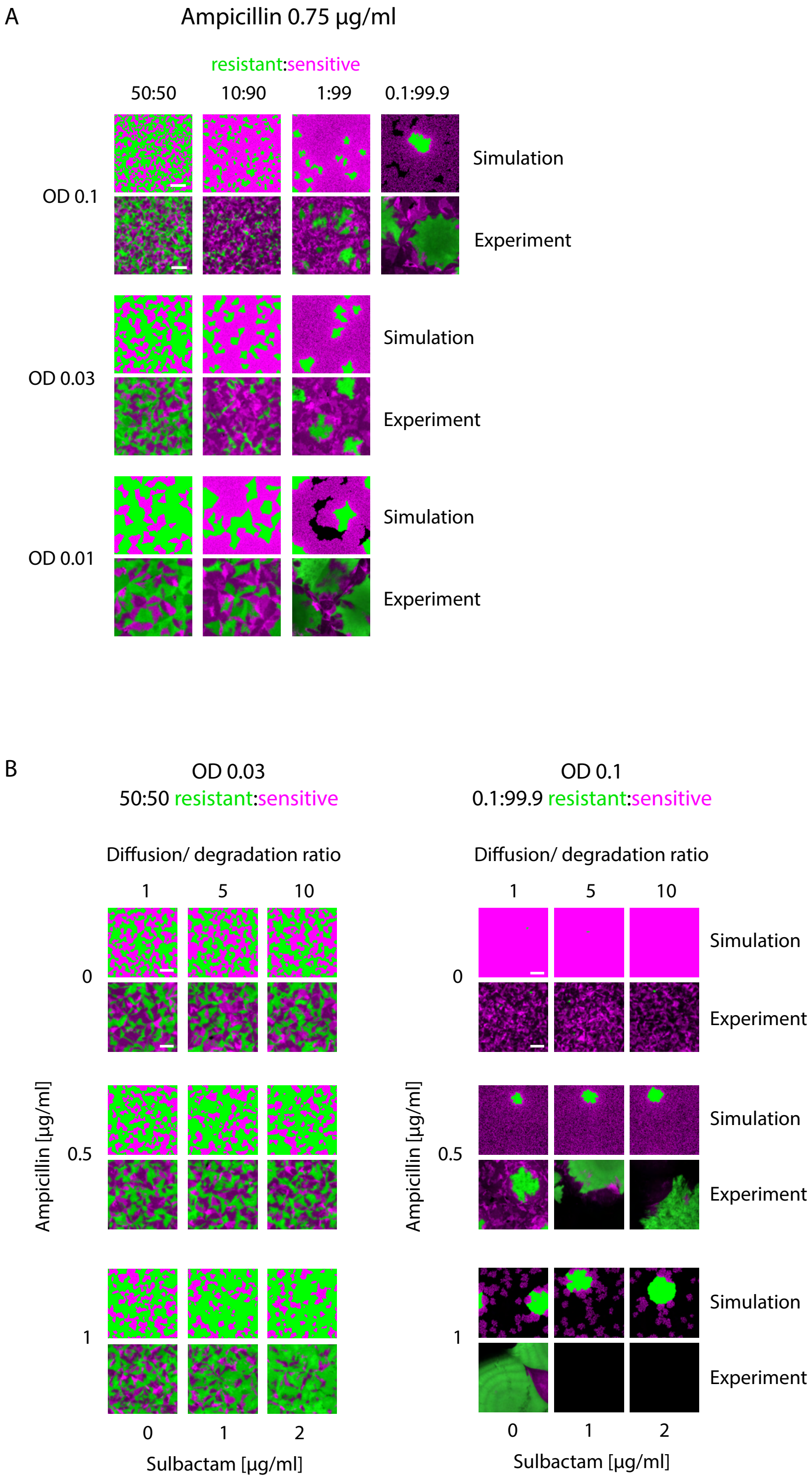


**Figure S15: Predictions of the biophysical model for a new ampicillin concentration and with addition of a β-lactamase inhibitor.** (A) Simulations (top) and experimental images for different initial densities and resistant:sensitive ratios at a previously untested ampicillin concentration of 0.75 μg/ml ampicillin. Resistant in green, sensitive in magenta, scale bars = 100 μm. (B) Simulations with degradation either every diffusion step (data same as Figures S10-12), every 5 diffusion steps, or every 10 diffusion steps to simulate inhibition of β-lactamase activity (top rows), compared to images of experimental colonies with 0, 1 or 2 μg/ml of the β-lactamase inhibitor sulbactam, for different ampicillin concentrations and two different initial colony compositions. Resistant green, sensitive magenta, scale bars = 100 μm.

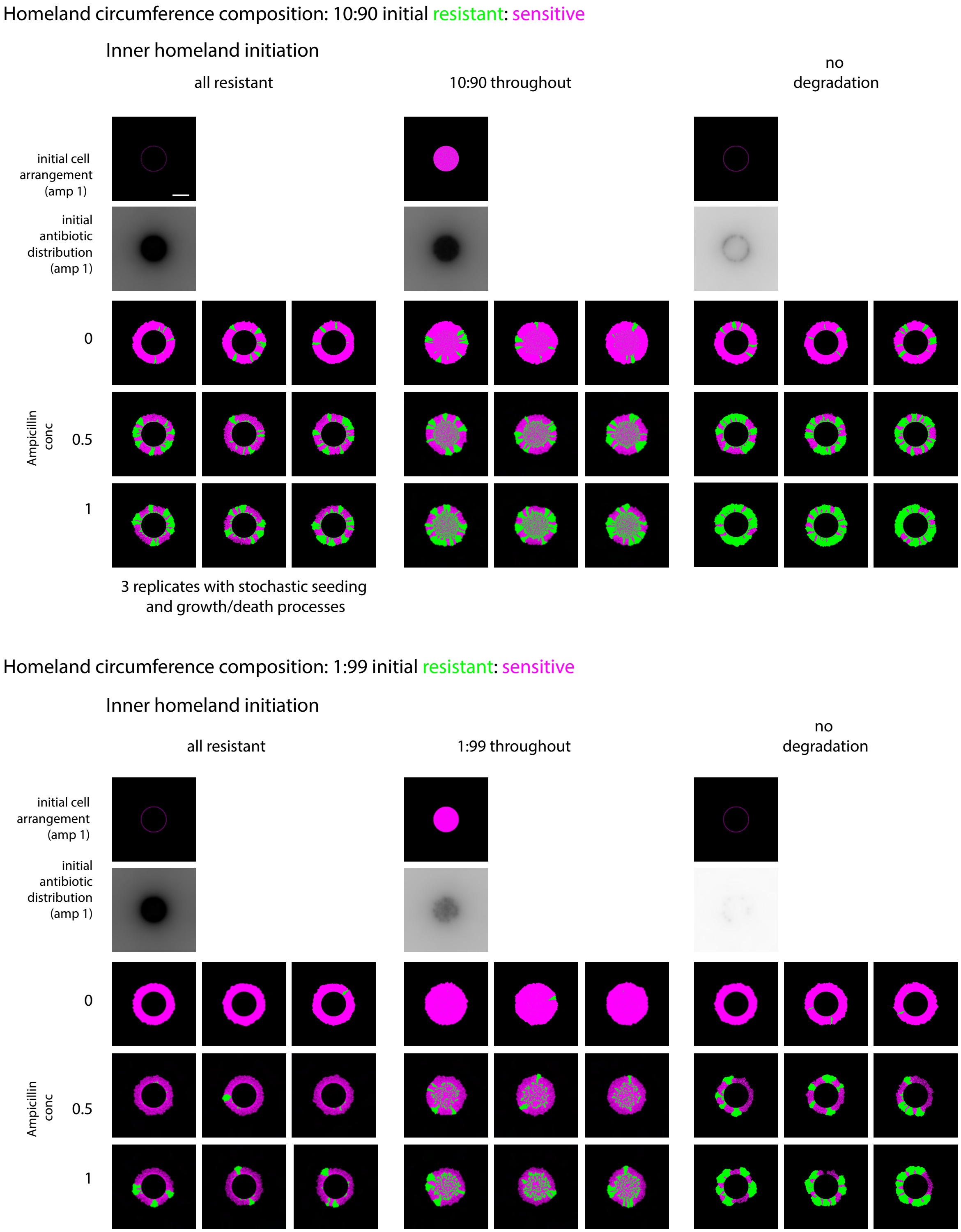


**S16: Effect of homeland-mediated antibiotic degradation on range expansion ecology is robust across replicates.** Replicates of simulations showing how degradation by resistant cells in the homeland could affect range expansion dynamics for two different initial resistant ratios at the expansion front and three different homeland compositions: all resistant/degrading, same ratio as the expansion front or non-degrading. In the all-resistant and no degradation cases, homeland cells do not contribute to growth, only to degradation of antibiotic if applicable. Initial cell arrangement and final range expansion patterns of resistant (green) and sensitive (magenta) bacteria for different global ampicillin concentrations are shown, as well as the antibiotic distribution at the initial time step. Scale bars = 100 µm.

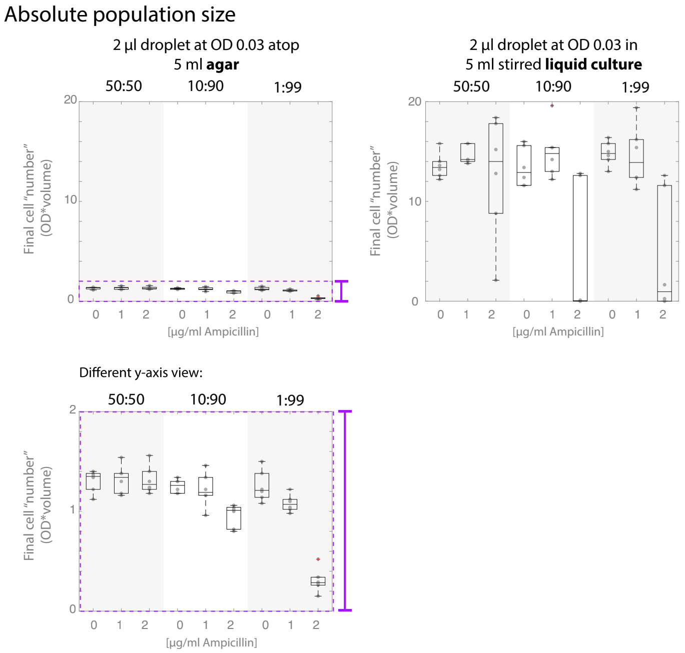


**S17: Total population sizes under ampicillin exposure in liquid culture versus agar colonies.** Comparison of total *E. faecalis* population sizes (calculated as measured optical density multiplied by volume of (diluted) culture, after ~18h of growth at 37˚C under ampicillin exposure, for populations grown on agar (left) versus in well-mixed liquid culture (right), at different initial resistant:sensitive ratios (indicated by shaded areas), all with an initial OD of 0.03. Bottom left panel shows same data from agar colonies as top left panel, with adjusted y-axis. Box plots with median and quartile lines and whiskers indicating data spread (apart from outliers) are shown, as well as individual data points (n = 6 for each condition). Outliers are defined as values more than 1.5 interquartile ranges from the edges of the box and shown as red '+'.

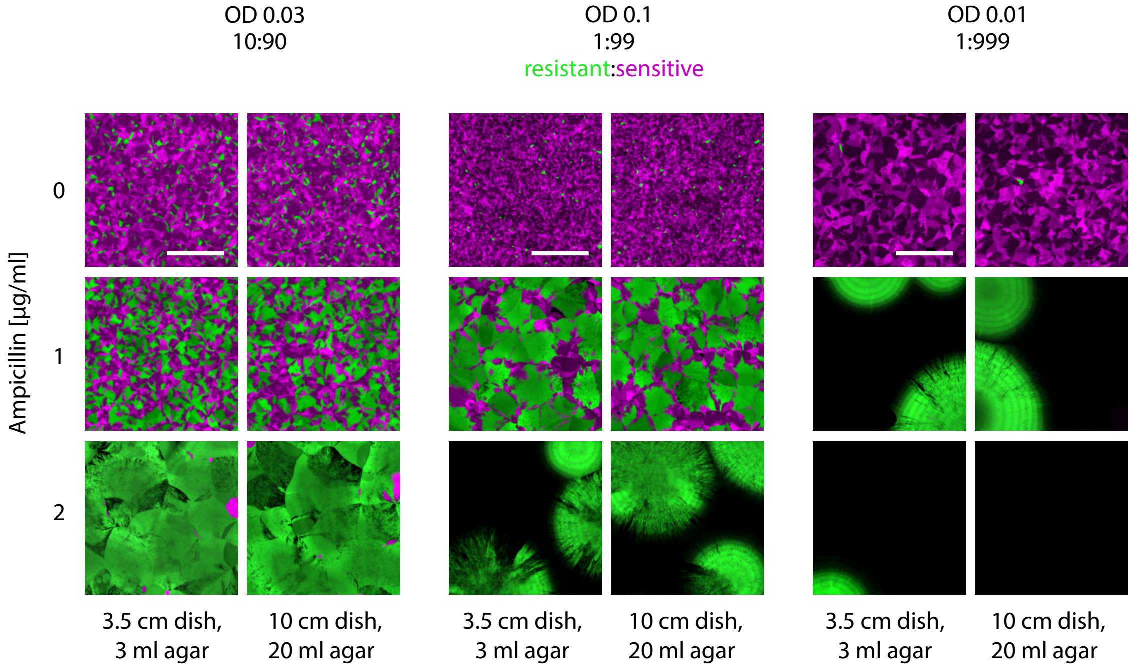


**Figure S18: Spatial patterns are robust to agar volume and height and thus to the global ampicillin reservoir.** Images of colonies across different initial conditions, grown either on 3ml agar on a 3.5 cm diameter petri dish, or 20 ml agar in a 10 cm diameter dish (all other experiments were conducted on 5ml agar in 3.5 cm dishes), to assess the relevance of height and volume of the agar containing ampicillin (and nutrients) on pattern formation. Resistant green, sensitive magenta, scale bars = 500 µm.

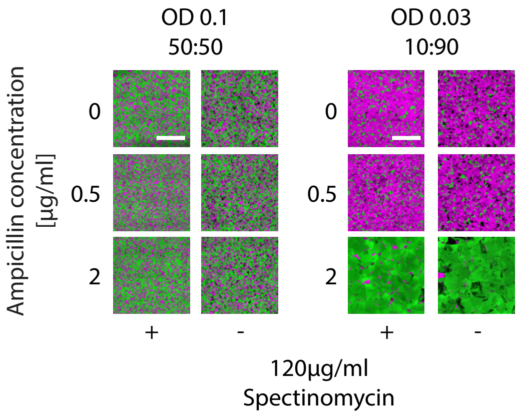


**Figure S19: Colonies grown without spectinomycin selection display more frequent fluorescence loss, but maintain similar spatial organization under ampicillin exposure as those with spectinomycin selection.** Confocal images of *E. faecalis* colonies grown on agar with (+)

our without (-) the antibiotic spectinomycin, which was otherwise used in all experiments to select for the pBSU101 plasmids carrying fluorescence and, if indicated, ampicillin resistance. In the presence of ampicillin, plasmids carrying the ampicillin resistance gene were presumably still selected for in the ampicillin-resistant strain (green), but not fluorescent plasmids without ampicillin resistance in the sensitive strain (magenta). Scale bars = 500 µm.

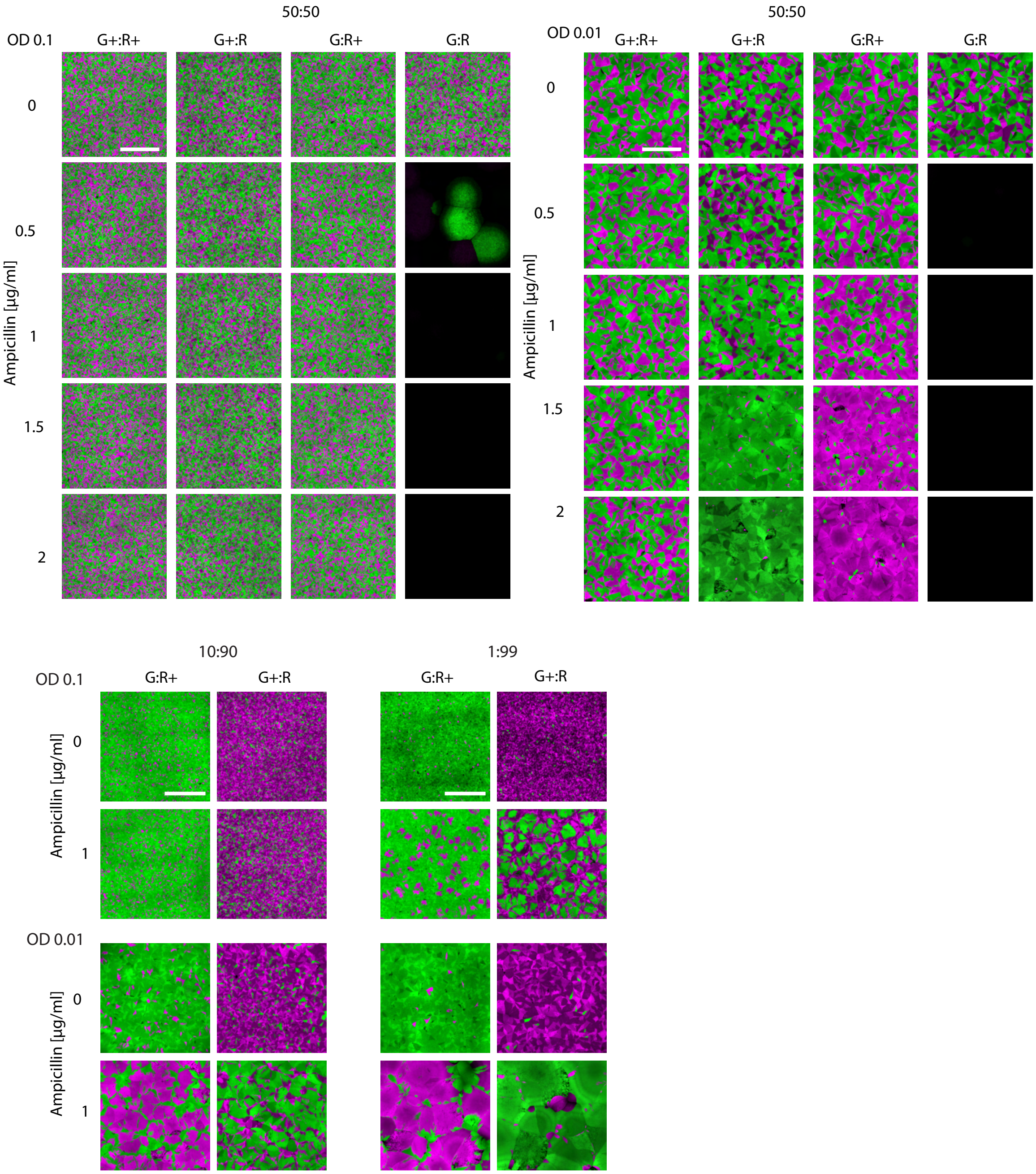


**Figure S20: *E. faecalis* strains with different fluorophores produce similar spatial organization.** Confocal images comparing homeland spatial patterns across ampicillin concentrations and community compositions for different combinations of pBSU101 plasmids carrying fluorescent proteins and, in some cases, the β-lactamase-encoding antibiotic resistance gene. G+ = mDasher with ampicillin resistance (green); R+ = mRudolph with ampicillin resistance (magenta); G = mDasher ampicillin sensitive (green); R = mRudolph ampicillin sensitive (magenta). Scale bars = 500 µm.

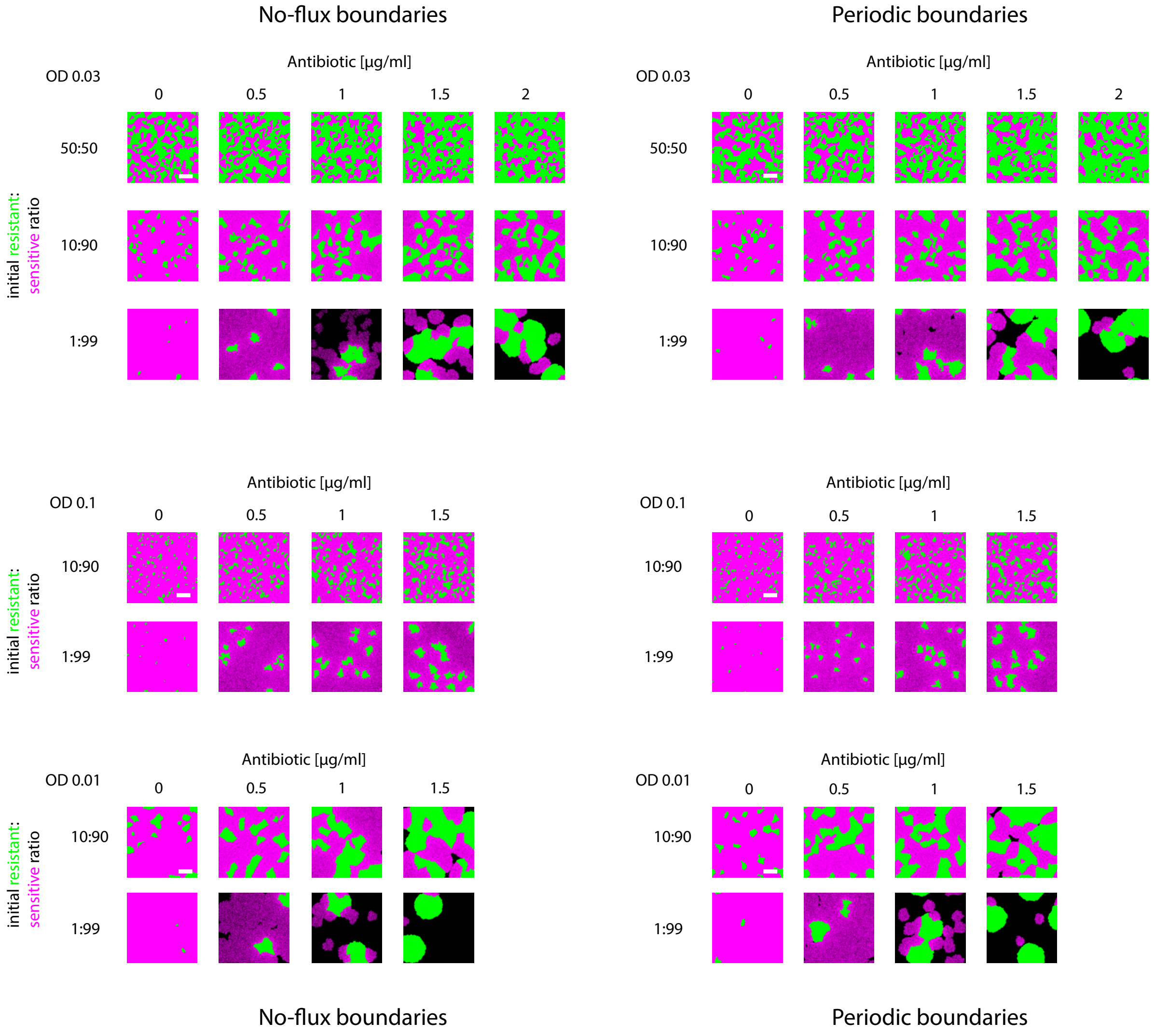


**Figure S21: Simulation results are robust to choice of boundary condition.** Simulations across various initial conditions, comparing a no-flux boundary condition (left column) in which division and diffusion do not surpass the boundary of the simulated area in x and y, to the periodic x-y boundaries (right column, same data as Figures S10-S12 shown here for direct comparison) used throughout the paper. Across most initial antibiotic and population conditions, simulations with periodic and no-flux boundary conditions yield similar patterns in terms of composition and length scale. Scale bars = 100 µm.

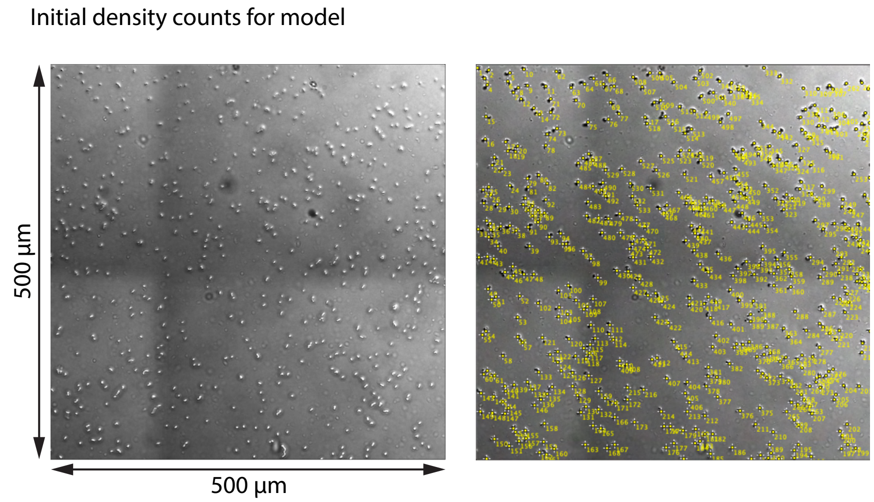


**Figure S22: Example of the experimental analysis used to determine initial seeding density for simulations.** Confocal image of an agar plate, at the center of the dried droplet containing the initially seeded population at OD 0.03, imaged within 45 min after seeding, without (left) and with (right) cell counts manually added in FIJI/ImageJ. Scale as indicated. These cell counts were used to inform initial cell seeding densities in the computational model.

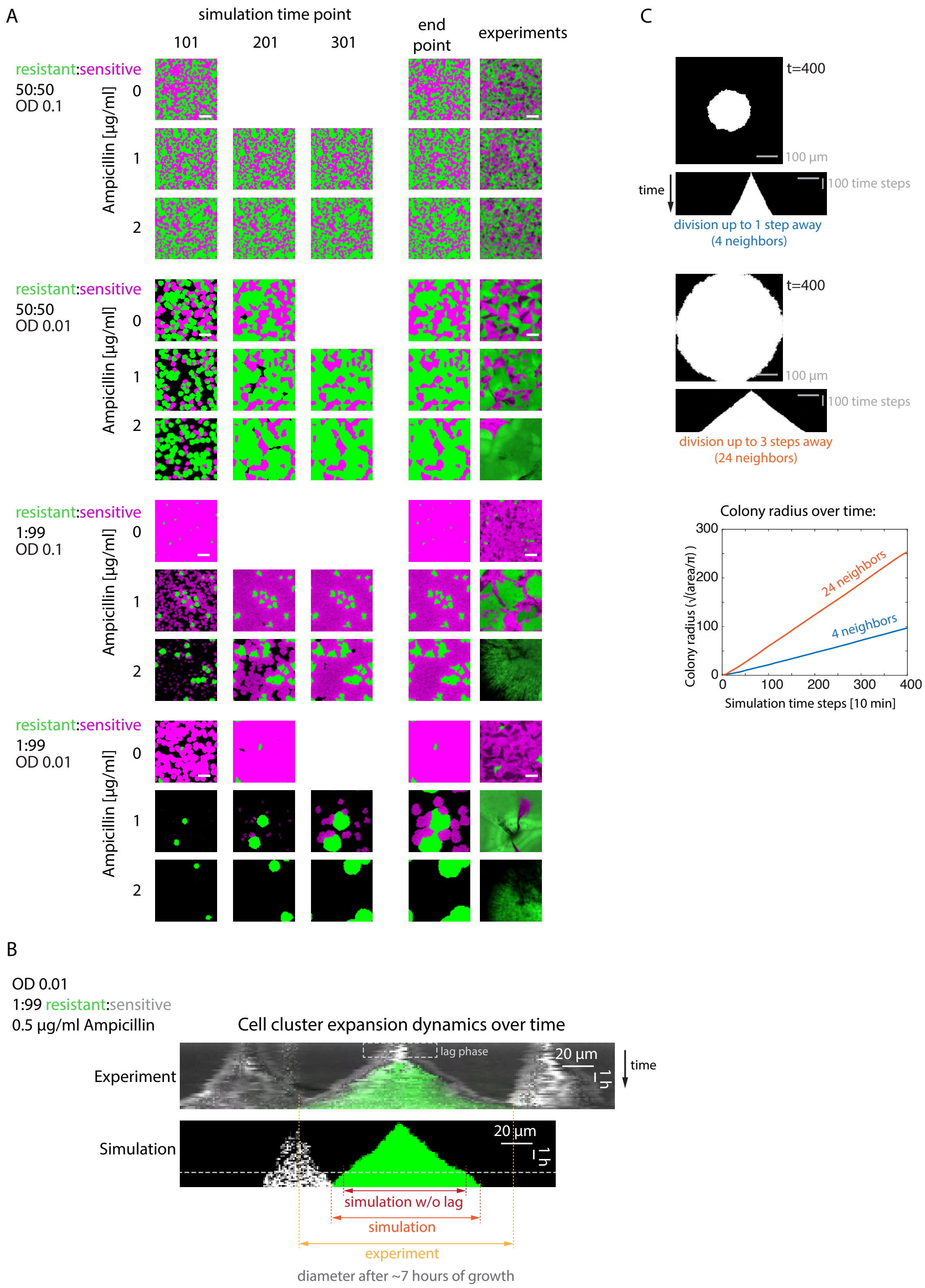


**Figure S23: Discrepancies between growth speed in simulations versus experiments were adjusted for by changing simulation run time and can be explained by local division rules.** (A)

Time series of simulations across different representative initial conditions (endpoints correspond to those shown in Figure 4 A) and corresponding experimental end point data. Resistant subpopulation is shown in green, sensitive subpopulation in magenta. While in cases with high initial densities, especially of resistant cells (which in our model grow independently of ampicillin concentration), full grid coverage similar to the experiment was reached after about 100 timepoints (equivalent to the experimental growth duration), other conditions took appreciably longer to grow to coverage similar to experimental data. A run time of 400 time points was chosen to approximately match overall coverage of experiments across conditions. This includes conditions in which the experiments did not exhibit full coverage of the agar surface. Scale bars = 100 µm. For cases without ampicillin in which the whole grid was filled before the full run time, simulations were ended early (since there is no death in absence of antibiotic and no growth is possible without space), so only those time points that were simulated are shown. (B) Comparison of the growth speed of experimental versus simulation clusters originated from a single cell over time. Images show kymographs (the same slice of the image (x direction) across timepoints (y direction) of a single resistant cluster (green), growing largely without competition (up until the final timepoints, when sensitive clusters (gray/white) start interfering). Both images show growth over about 7 hours. While the diameter of the simulation cluster (bottom) increases at a linear rate, the experimental cluster initially does not grow appreciably (lag phase), but then expands rapidly. Its growth rate appears to increase rapidly toward the end of the acquisition, causing the cluster to become larger than the simulation cluster, especially when the non-growth phase is adjusted for ("simulation w/o lag"). (C) Simulation data testing how the restriction of growth to open cluster surfaces affects effective growth rate (in the absence of antibiotic). In our model, the appearance of a new cell is only allowed in an open space that is nearest neighbor to the dividing cell (4 neighbors). This effectively only allows cells directly at the surface of a cluster to divide, similarly to classic models like the Eden growth model, leading to a slow linear increase in colony radius. If, in contrast, we allow new cells to appear up to three grid steps away from a dividing cell (24 neighbors), this effectively allows cells one or two rows behind the surface to divide (with lower probability), thus creating a wider active layer. This simulation grows linearly, albeit at a faster rate, even though the baseline division probabilities (before accounting for availability of space) are identical for both cases. Data is displayed as final timepoint grid and kymograph of time evolution for both models (top), and a quantification of colony radius change over time (bottom).

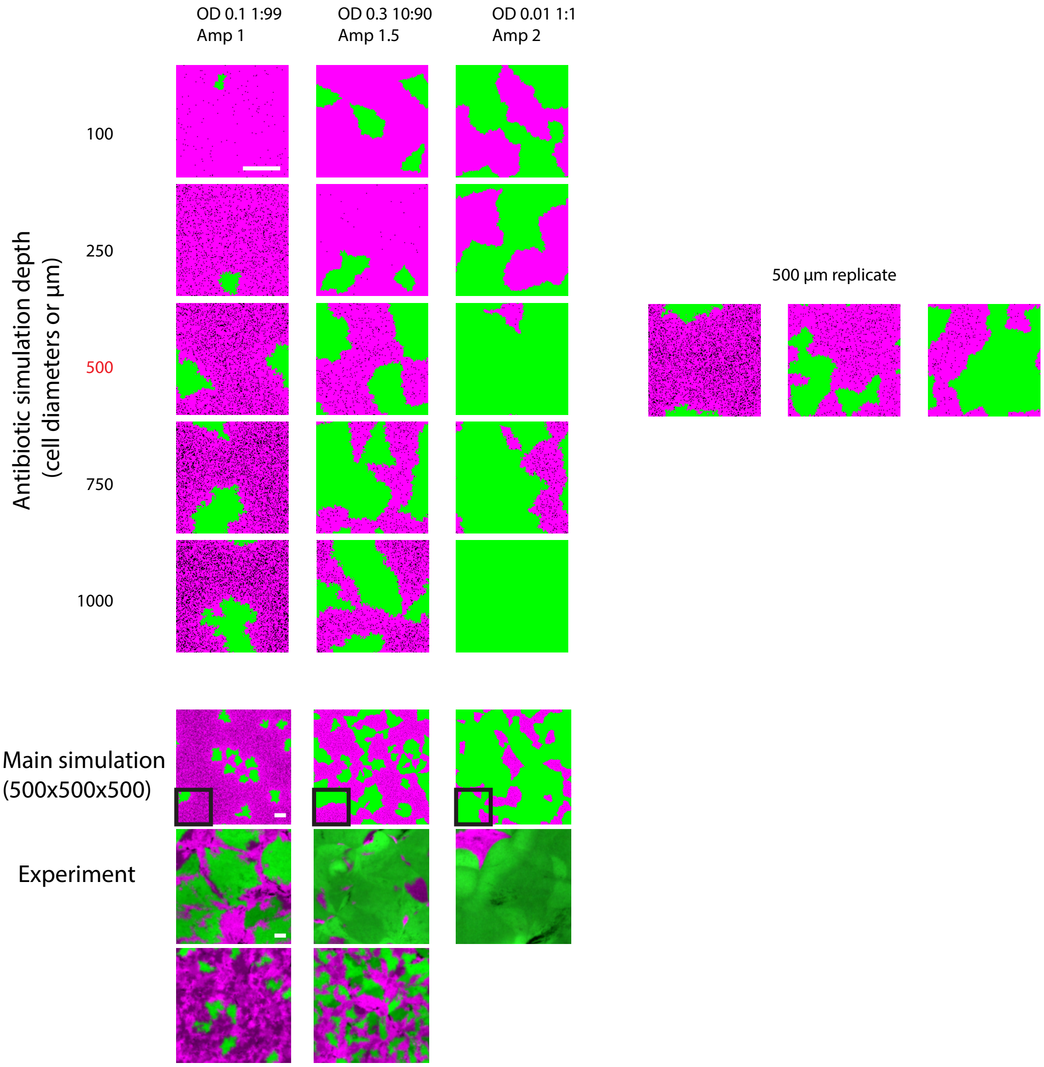


**Figure S24: Changing the depth of the antibiotic simulation only modestly alters outcomes beyond 500 µm.** Simulations on 100 µm x100 µm cell grids, varying the depth of the antibiotic between 100 and 2000 µm (500 µm is the depth of the main simulation dataset, shown below with experimental data for comparison). A replicate simulation at 500 µm depth is shown on the right, to assess simulation variability under these conditions. Black squares on "main simulation" show the relative size of simulations above it. A smaller grid size was used due to computational limitations for these simulations. Scale bars = 50 µm.

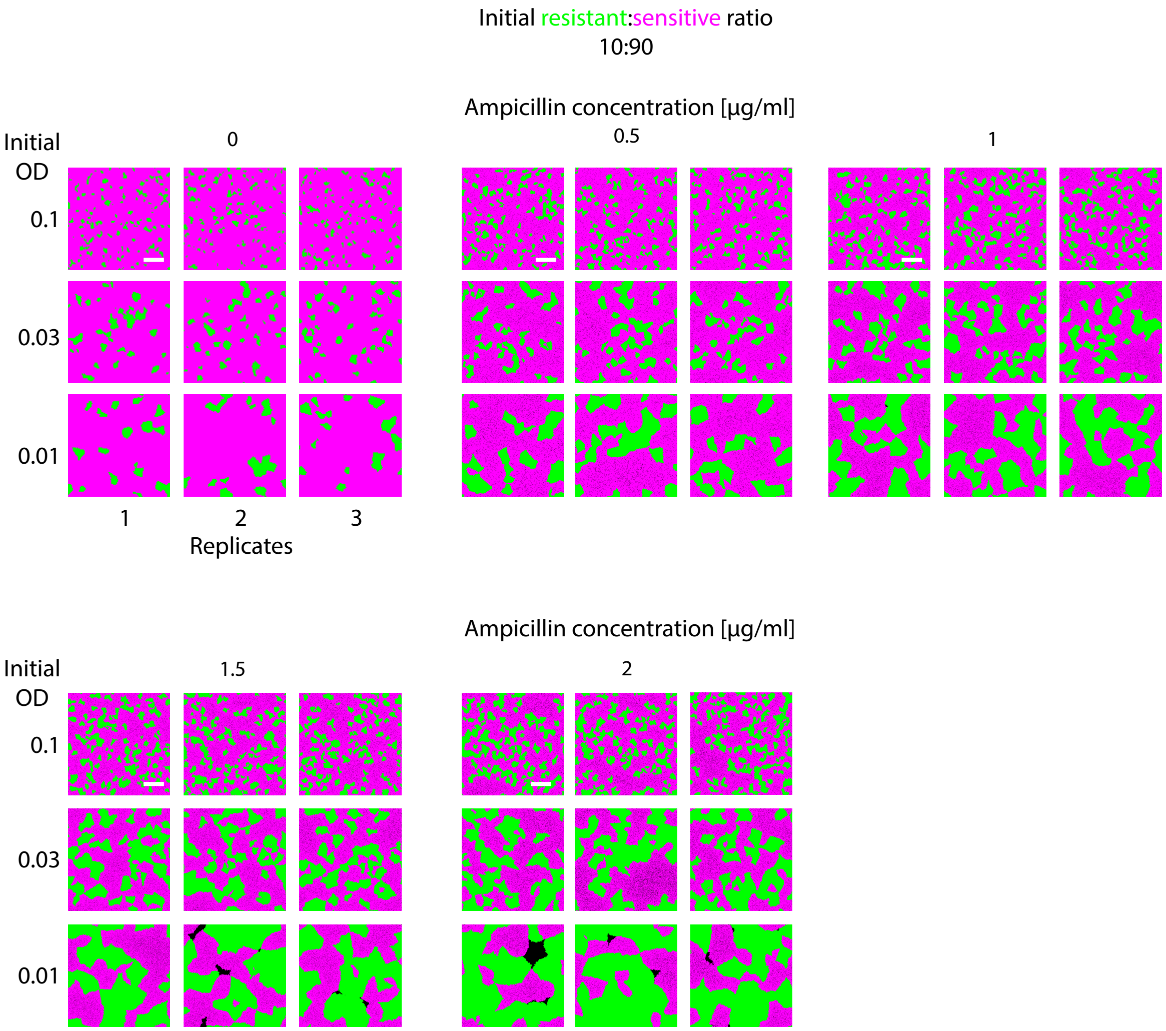


**Figure S25: Spatial composition and scale is robust across simulation replicates with stochastic seeding, growth, and death.** Final cell grids of three replicate 500 µm x 500 µm x 500 µm simulations (shown as neighboring columns) with identical initial cell numbers and ratios, but independent stochastic initial positions and growth and death rates, tested across different initial densities and global ampicillin concentrations, at an initial resistant(green):sensitive(magenta) ratio of 10:90. While exact distributions varied between replicates, overall composition and pattern size were similar. Scale bars = 100 µm.

Supplemental Movie Legends:

**Movie S1: Spatial bacterial simulation dynamics at relatively high resistant fraction.** Resistant (green) and sensitive (magenta) bacterial growth and death dynamics over all simulation time points at an initial density equivalent to OD 0.03 in the initial droplet (see Methods for estimations of seeding density), 10:90 resistant:sensitive initial ratio, 1.5µg/ml global ampicillin concentration. Every 5$^{th}$ time point is shown, 7 frames per second, one time point equates around 10 minutes relative to doubling time and diffusion rates. Final timepoint labeled "401" is actually the last simulation time step (400). Scale bar = 100 µm.

**Movie S2: Spatial antibiotic simulation dynamics at relatively high resistant fraction.** Antibiotic concentration (greyscale, black = 0µg/ml, white = 1µg/ml or higher) over all simulation time points at an initial density equivalent to OD 0.03 in the initial droplet (see Methods for estimations of seeding density), 10:90 resistant:sensitive initial ratio, 1.5µg/ml global ampicillin concentration. Every 5$^{th}$ time point is shown, 7 frames per second, one time point equates around 10 minutes relative to doubling time and diffusion rates. Final timepoint labeled "401" is actually the last simulation time step (400). Scale bar = 100 µm.

**Movie S3: Spatial bacterial simulation dynamics at low antibiotic concentration and low resistant fraction.** Resistant (green) and sensitive (magenta) bacterial growth and death dynamics over all simulation time points at an initial density equivalent to OD 0.01 in the initial droplet (see Methods for estimations of seeding density), 0.1:99.9 resistant:sensitive initial ratio, 0.5 µg/ml global ampicillin concentration . Every 5$^{th}$ time point is shown, 7 frames per second, one time point equates around 10 minutes relative to doubling time and diffusion rates. Final timepoint labeled "401" is actually the last simulation time step (400). Scale bar = 100 µm.

**Movie S4: Spatial antibiotic simulation dynamics at low antibiotic concentration and low resistant fraction.** Antibiotic concentration (greyscale, black = 0µg/ml, white = 1µg/ml or higher) over all simulation time points at an initial density equivalent to OD 0.01 in the initial droplet (see Methods for estimations of seeding density), 0.1:99.9 resistant:sensitive initial ratio, 0.5µg/ml global ampicillin concentration. Every 5$^{th}$ time point is shown, 7 frames per second, one time point equates around 10 minutes relative to doubling time and diffusion rates. Final timepoint labeled "401" is actually the last simulation time step (400).Scale bar = 100 µm.

**Movie S5: Spatial bacterial simulation dynamics at relatively high antibiotic concentration and low resistant fraction.** Resistant (green) and sensitive (magenta) bacterial growth and death dynamics over all simulation time points at an initial density equivalent to OD 0.03 in the initial droplet (see Methods for estimations of seeding density), 0.1:99.9 resistant:sensitive initial ratio, 1 µg/ml global ampicillin concentration. Every 5$^{th}$ time point is shown, 7 frames per second, one time point equates around 10 minutes relative to doubling time and diffusion rates. Final timepoint labeled "401" is actually the last simulation time step (400). Scale bar = 100 µm.

**Movie S6: Spatial antibiotic simulation dynamics at relatively high antibiotic concentration and low resistant fraction.** Antibiotic concentration (greyscale, black = 0µg/ml, white = 1µg/ml or higher) over all simulation time points at an initial density equivalent to OD 0.03 in the initial droplet (see Methods for estimations of seeding density), 0.1:99.9 resistant:sensitive initial ratio, 1µg/ml global ampicillin concentration. Every 5th time point is shown, 7 frames per second, one time point equates around 10 minutes relative to doubling time and diffusion rates. Final timepoint labeled “401” is actually the last simulation time step (400). Scale bar = 100 µm.
